# A Framework for Identifying Determinants of Epigenomic State: Decoding Variable CTCF Occupancy and Contact Frequency

**DOI:** 10.64898/2026.11.25.769053

**Authors:** Yunxiao Yang, Michael A. Beer

## Abstract

CTCF binding is essential for delineating topologically associated domains, constraining enhancer-promoter interactions, and controlling gene expression. Many CTCF peaks are bound in all cell-types, but the mechanisms regulating cell-type-specific CTCF occupancy are not fully understood. Here, we present a general framework to comprehensively discover factors driving variable epigenomic state and TF occupancy. We apply this framework to CTCF ChIP-seq binding across hundreds of cell/tissues, and we identify five distinct mechanisms regulating CTCF occupancy. In addition to known mechanisms of DNA methylation and cell-type-specific TF co-binding, we find that the largest set of variable-occupancy sites is modulated by CTCF expression levels. Variable peaks have weaker-affinity, and their occupancy is modulated by either higher CTCF levels or flanking TFs. We evaluate the predictive power on CTCF occupancy in held-out samples. Finally, we show that variable CTCF binding predicts differential contact frequency, chromatin loop structure, and upregulation of gene expression.

## Introduction

Transcriptional regulation is essential for controlling cell-state transitions and the maintenance of cellular homeostasis. This process is achieved through interaction between *trans*-regulatory elements and *cis*-regulatory elements (CREs). The former are transcription factors (TFs), which are proteins with DNA-binding domains, and the latter are noncoding genomic regions bound by transcription factors. TF binding to CREs can alter the transcription rate of nearby genes, leading to phenotypic changes. Therefore, characterizing and understanding the mechanisms driving differential TF binding is key to deciphering changes in the transcriptome.

Current studies of differential TF occupancy often focus on specific postulated mechanisms. While this hypothesis-driven approach has provided invaluable insights, and common mechanisms like changes in DNA methylation can explain changes in TF binding, it may cause other promising explanations to be missed. Without considering all mechanisms together, the current approach also falls short in quantifying the fraction of TF binding sites (TFBS) regulated by each mechanism. Therefore, we propose a more general analysis framework which is not specific to a particular mechanistic explanation. We then applied this approach to elucidate the factors driving cell-type-specific CTCF binding, given its importance in genome organization, and the large number of available ChIP-seq datasets for CTCF. Additionally, the strong conservation of CTCF-binding sites across mammals^1^ supports their important regulatory role.

CTCF, a ubiquitously expressed TF, blocks DNA extrusion of the cohesin complex to create chromatin loops and form topologically associated domains (TADs).^2–5^ CREs and promoters within the same TAD interact more frequently, and these interactions control transcription levels for genes in the TAD.^6^ Therefore, differential CTCF binding can modulate gene expression by adding or removing TAD boundaries,^7^ which thereby modulate CRE-promoter interactions. Thus, knowing why CTCF binds variably is essential to understanding cell-type-specific transcriptional regulation.

It has long been known that DNA CpG methylation can inhibit TF binding, and it has previously been reported that differential methylation regulates 40% of variable CTCF binding.^8^ Some CpG positions directly within CTCF binding sites can be methylated, hindering CTCF-DNA interaction.^9–11^ Other studies have linked hypermethylation in embryonic development to CTCF binding loss.^12^ However, methyltransferase knockout strengthens occupancy at only 7% of sites^11,12^ and reactivated less than 2% of previously methylated sites,^11^ suggesting methylation is not the primary driver of changes in CTCF binding. Further, only 10% of CTCF binding sites have a CpG,^13^ so methylation is unlikely to explain variation at most sites. While others recently have promoted cell-type-specific TFs as regulators of CTCF binding,^14–18^ it remains unclear what fraction of CTCF peaks are regulated through this mechanism.

To address this gap, we developed an analysis framework. We began by uniformly processing hundreds of CTCF ChIP-seq ENCODE datasets and constructing a list of consensus peaks from tissue and cell-line samples (**Fig. 1A**). We then clustered peaks by shared occupancy patterns across datasets. After identifying distinct classes of co-variable peaks, we explained their variability by cell-type-specific epigenetic and non-epigenetic mechanisms. While some variable peaks are associated with DNA methylation or cell-type-specific TFs, surprisingly, more were modulated by CTCF expression. We observed that peaks regulated by different mechanisms also had different binding-site affinities, supporting a model in which TFBS strength and flanking sequence context contribute to the dominant regulatory mechanisms. To show that our framework quantified most of the variability, we trained a simple regression model to predict CTCF occupancy on held-out samples. Finally, we show that differential contact frequency is predicted by differential CTCF binding and that gene upregulation is often accompanied by stronger and more stable chromatin loops.

**Figure 1:**
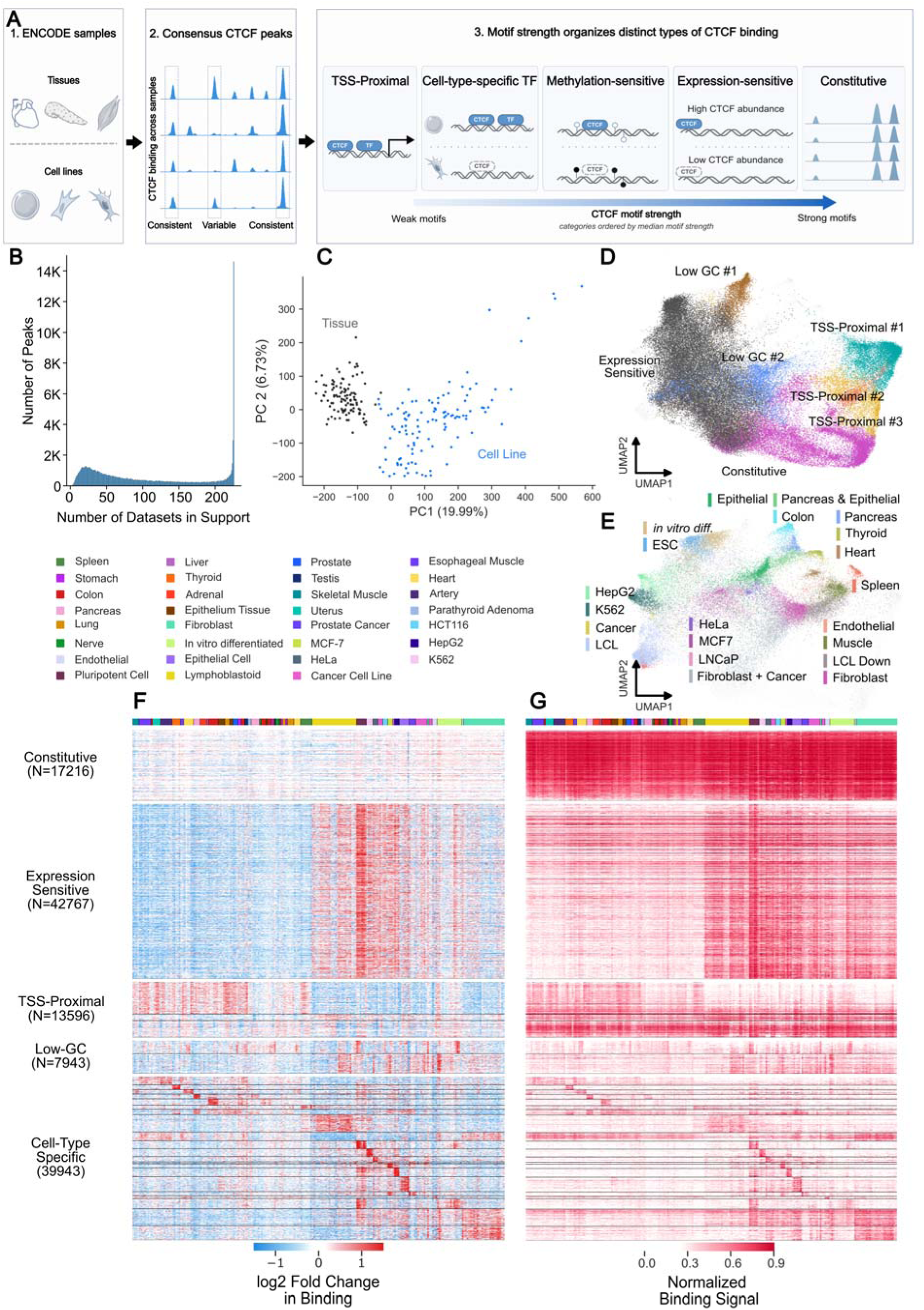
Variable modes of CTCF binding across cell/tissue types. (**A**) A schematic of the workflow to understand mechanisms controlling variable CTCF occupancy. Starting with a large ENCODE data collection of CTCF ChIP-seq experiments, we built a consensus peak list, clustered peaks based on distinct binding variations across samples, and explained these variations with different mechanisms. (**B**) The number of datasets supporting consensus CTCF peaks. Support was defined as the number of experiments in which peaks passed the statistical threshold. (**C**) Principal component analysis (PCA) of CTCF ChIP-seq datasets using the consensus binding matrix. Experiments assayed on tissue samples and on cell-line samples were highlighted in gray and blue, respectively. UMAP embedding of consensus peaks, with constitutive, expression-sensitive, TSS-proximal, and low-GC peaks highlighted in (**D**) and cell-type-specific peaks highlighted in (**E**). Heatmaps showing (**F**) variation in normalized CTCF occupancy and (**G**) normalized CTCF occupancy for peaks across biological samples. Peaks were grouped by annotated clusters, and samples were hierarchically clustered in both heatmaps. Cluster categories were annotated on the left. Variation in (**F**) was computed as the log_2_ fold change in binding relative to the average binding across all datasets. For visualization, binding in (**G**) was scaled to the range [0, 1].

**Extended Data Figure 1:**
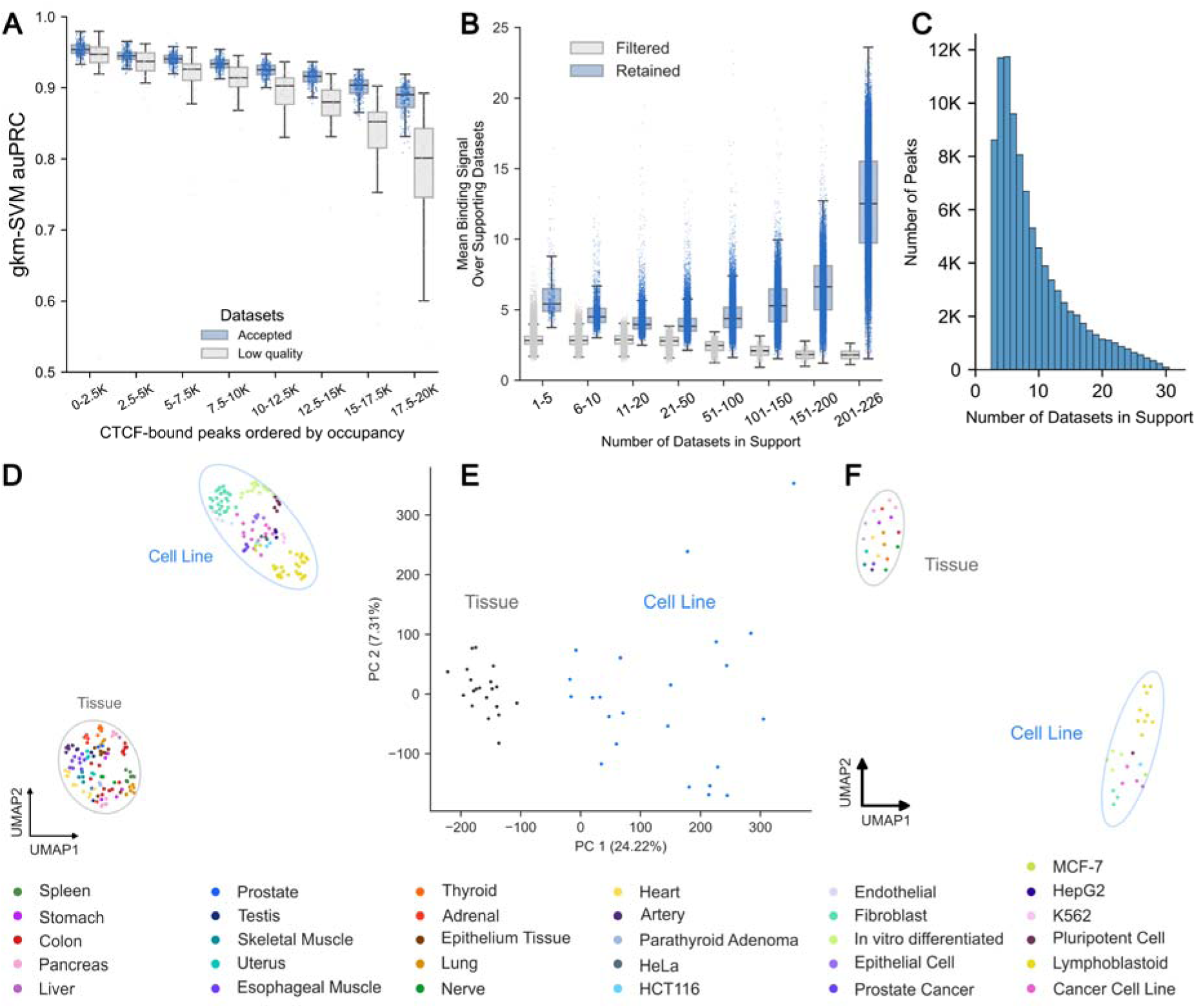
Conduct quality control and characterize consensus CTCF peaks. (**A**) gkm-SVM was used to classify the strongest CTCF peaks from random genomic background. Cross-validation auPRC scores were used to quality-control CTCF ChIP-seq datasets. Each point in the box plot represented 2500 peaks from one dataset, ranked by binding signal. A lower auPRC suggested a poorer signal-to-noise ratio. High-quality datasets generally had higher auPRC than low-quality experiments, which showed a much sharper decline in auPRC as peak binding strength decreased. (**B**) Peaks in the consensus list had much higher average binding than those removed. Each point in the box plot denoted one peak. The distribution of average binding was stratified by the number of datasets in support. (**C**) The number of datasets supporting consensus peaks, constructed from JUN, JUNB, and JUND. Supports were defined as the number of datasets where peaks had statistically significant signals. (**D**) UMAP embedding of CTCF ChIP-seq datasets. Point colors represented the sample characterized by the dataset. For a subset of CTCF ChIP-seq datasets generated by the same lab with the same antibody batch, low-dimensional embeddings were recalculated using (**E**) PCA and (**F**) UMAP. As with the full set, datasets characterizing tissue samples were separated from those characterizing cell-line samples along the first principal component, and datasets assayed on similar biological samples were clustered in UMAP space.

## Results

### Analysis framework to find peaks with distinct binding patterns

We first assessed the quality of the CTCF ChIP-seq experiments using gkm-SVM predictive performance, as described in gkmQC (**Methods**).^19–22^ This approach uses the sequence-based machine learning model gkm-SVM. Actual peaks are driven by sequence-features under the peaks, while noise is not predictable from underlying sequence, and importantly, this varies consistently with measured peak strength. Most datasets are well predicted by the gkm-SVM model, but some outlier datasets with lower model performance likely result from poor signal-to-noise ratios which cause CTCF motif-absent regions to be called as peaks (**Extended Data Fig. 1A**). Those few low-scoring outlier experiments we deemed low quality and removed them from subsequent analyses. Datasets that characterized pooled samples or had weak cross-replicate correlation were also removed. In total 76 of 302 datasets were removed. This might seem overly conservative, but since we are specifically trying to model variation in peak strength across samples, it is critical to remove noise as a source of peak variability.

To create a consensus peak list, we used a two-step filtering procedure (**Methods**). We first aggregated peaks that met significance thresholds across all high-quality ChIP-seq experiments. Then we pruned weaker binding strength peaks, using actual binding signals in each experiment, as weakly bound peaks could appear statistically significant in high-read-depth experiments (**Extended Data Fig. 1B**). This procedure resulted in 126K high-confidence consensus CTCF peaks (**Supp. Table 1**). Some of these are constitutively detected across datasets; however, the majority show variable occupancy, consistent with previous observations (**Fig. 1B**).^8,11,23^ This is in sharp contrast with other non-CTCF TFs, which show no evidence of ubiquitous binding sites, and whose occupancy is mostly controlled by combinatorial interactions with cell-type specific co-factors,^21^ as shown for JUN family proteins in **Extended Data Fig. 1C**.

After constructing the binding matrix, we next characterized how CTCF binding profiles varied across biological samples with dimensionality reduction, first with PCA. Most variation, along the first principal component, separated datasets assayed on tissue samples from those assayed on cell-line samples, explaining 20% of the total binding variance (**Fig. 1C**). As our later analysis will demonstrate, this cell-line vs tissue variance is driven by a specific subset of peaks which respond to CTCF expression, and this PCA component is correlated with CTCF gene expression across samples (R=0.64), but the fact that the strongest binding sites do not vary with expression implies that this is not a technical artifact. In the UMAP space, datasets also primarily clustered by cell-line vs tissue, but on finer scales clustered by biological sample type (e.g. LCL vs. fibroblast, **Extended Data Fig. 1D**). This pattern persists even when we subset the datasets to control for differences in source lab and specific antibody choice, suggesting that these observations are not due to these potential technical artifacts (**Extended Data Fig. 1E-F**).

We then clustered peaks to identify peaks with shared binding variability across biological samples (**Methods**).^24,25^ This approach yielded 28 fine-grained clusters with distinct occupancy profiles (**Fig. 1D-E**). Based on their epigenomic and non-epigenomic characteristics, we grouped them into six broad categories: constitutive, expression-sensitive, transcription start site (TSS)-proximal, methylation-sensitive, low-GC-content, and cell-type-specific, with the methylation-sensitive cluster as a subclass of the TSS-proximal category (**Supp. Table 2**, **Fig. 1F-G**). Next, we discuss each category and the mechanisms that regulate its binding in depth, except for the low-GC group, which had the weakest CTCF binding and no clear mechanisms explaining its binding variation.

### Constitutive CTCF peaks have high-affinity CTCF motifs

The first category of peaks is constitutively bound across all biological samples and have the strongest binding signal and CTCF motif strength. CTCF motif score approximates the binding free energy of the CTCF-DNA interaction, with higher scores indicating more favorable interactions and more stable binding.^26,27^ We scanned each 300bp peak sequence for its strongest CTCF binding site. Each peak was thus assigned a motif score that reflected its binding site’s similarity to the CTCF motif from JASPAR (**Methods**).^28,29^ **Fig. 2A** shows the fraction of peaks from each category binned by CTCF motif strength and also by ChIP-seq binding signal. Most peaks with high motif-scores and strongest binding signals are constitutive. Peaks in the other categories, such as cell-type-specific or TSS-proximal, had lower motif scores and weaker ChIP-seq binding signal.

**Figure 2:**
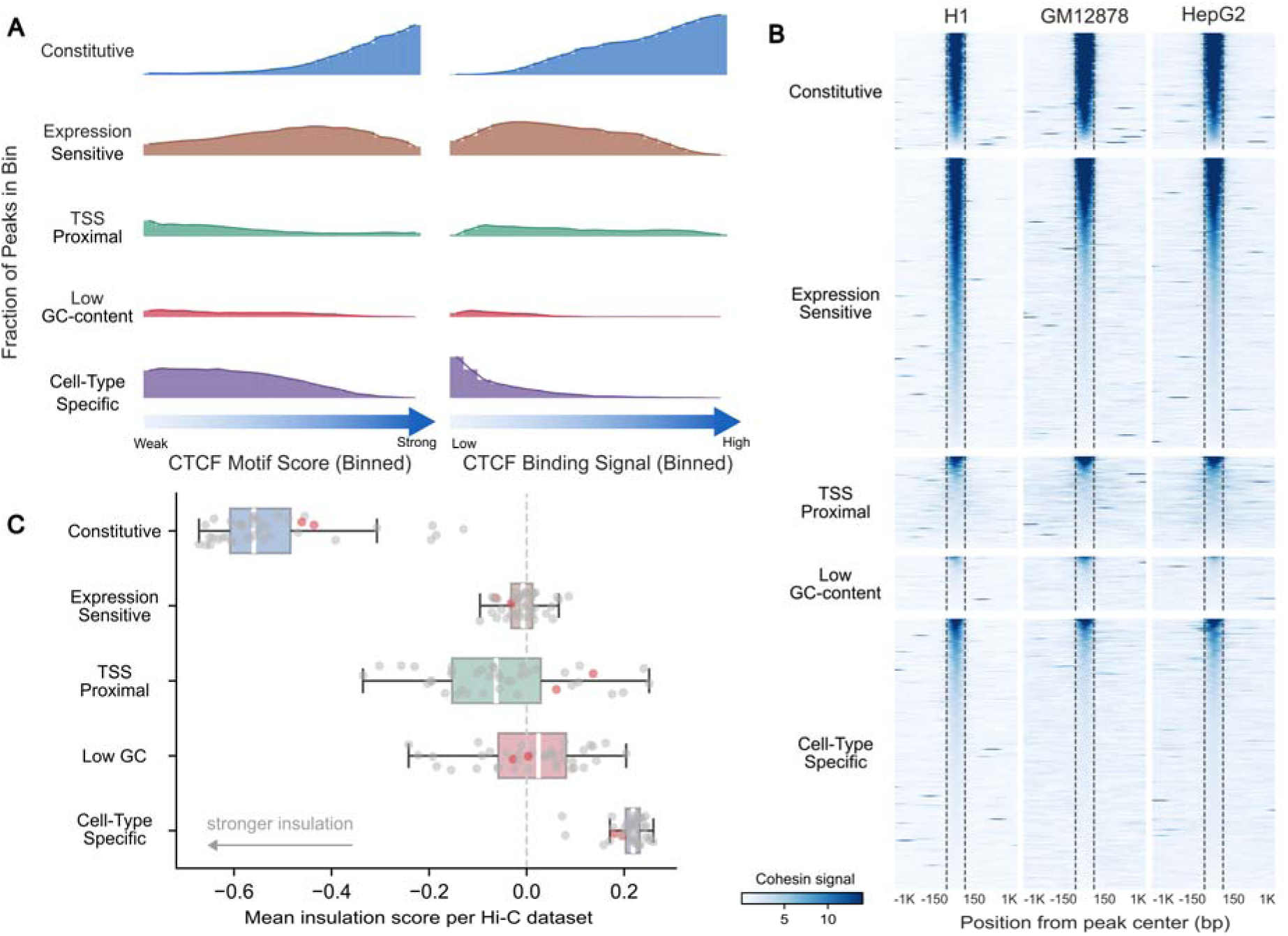
Constitutive CTCF peaks had the strongest CTCF occupancy, cohesin occupancy, and insulation strength. (**A**) Fraction of consensus peaks, binned by CTCF motif score or average binding strength, belonging to each category. Each peak’s motif score was defined as the highest score assigned to a binding site via motif scanning. Constitutive peaks accounted for the largest fraction of peaks in the highest bins for both motif score and binding strength. (**B**) Tornado plot showing the strength of cohesin subunit (SMC3 and RAD21) binding at the consensus peaks. Per-base-pair binding was visualized for regions ±1 kb around the peak center. Peaks were stratified by category, and within each category, peaks were sorted by the average cohesin occupancy. (**C**) Mean insulation score across peak categories. Each point represented one Hi-C dataset. HepG2 and GM12878 Hi-C were highlighted in red. A more negative score entailed a stronger insulation effect.

CTCF facilitates genomic organization by blocking the cohesin complex,^30^ so we next assess each peak category by its ability to retain cohesin. Across three separate cell lines, the constitutive peaks showed the strongest and most consistent cohesin binding (SMC3 and RAD21 ChIP-seq, **Fig. 2B**), consistent with their role as stable chromatin loop anchors, and as previously described over a more limited set of cell types.^31,32^ While some peaks in the expression-sensitive category also exhibited strong cohesin occupancy, the fraction of such peaks was smaller and more variable across cell types, ranging from about 50% in H1 to about 25% in cancer cell lines. Peaks in other categories had weaker cohesin binding.

Stronger cohesin binding at constitutive CTCF sites also translates to stronger TAD insulation, whereby regions between two loop-forming CTCF peaks interact less frequently than regions outside the loop. This decrease in interaction frequency can be quantified with insulation scores.^33,34^ Across different Hi-C samples, constitutive peaks showed the strongest insulation effect (**Methods**, **Fig. 2C**). Other categories exhibited weaker insulation, and cell-type-specific peaks were, on average, the weakest insulators. Overall, cohesin enrichment and insulation scores both supported constitutive peaks as the most robust TAD anchors.

### Moderate-strength CTCF binding sites are more sensitive to CTCF expression differences

#### A large set of CTCF peaks showed expression sensitivity

We observed a large class of variable CTCF peaks that were more strongly bound in cell lines compared to tissues but were not explained by simple sequence properties. We found that this set of CTCF peaks had their occupancy strongly positively correlated with CTCF gene expression across datasets (**Methods**; **Fig. 3A**). Peaks in this cluster have a median Spearman correlation of 0.43 with expression, and over 98% of peaks are positively correlated with CTCF expression (**Fig. 3B**). We thus annotated this cluster as “expression-sensitive,” and it turns out to be the largest cluster, accounting for 35% of the consensus peak list (**Extended Data Fig. 5F**). In contrast, all other clusters showed a much weaker correlation, ranging from −0.13 to 0.18. The mean binding signal averaged over all peaks in the expression-sensitive cluster is correlated with expression at 0.55; meanwhile, peaks in the constitutive cluster remained uninfluenced by CTCF expression (**Extended Data Fig. 2A**). While some cell-type-specific clusters exhibited moderate correlations, these trends were mostly driven by samples in which the peaks were enriched, since the average occupancy was largely stable outside of those samples.

**Figure 3:**
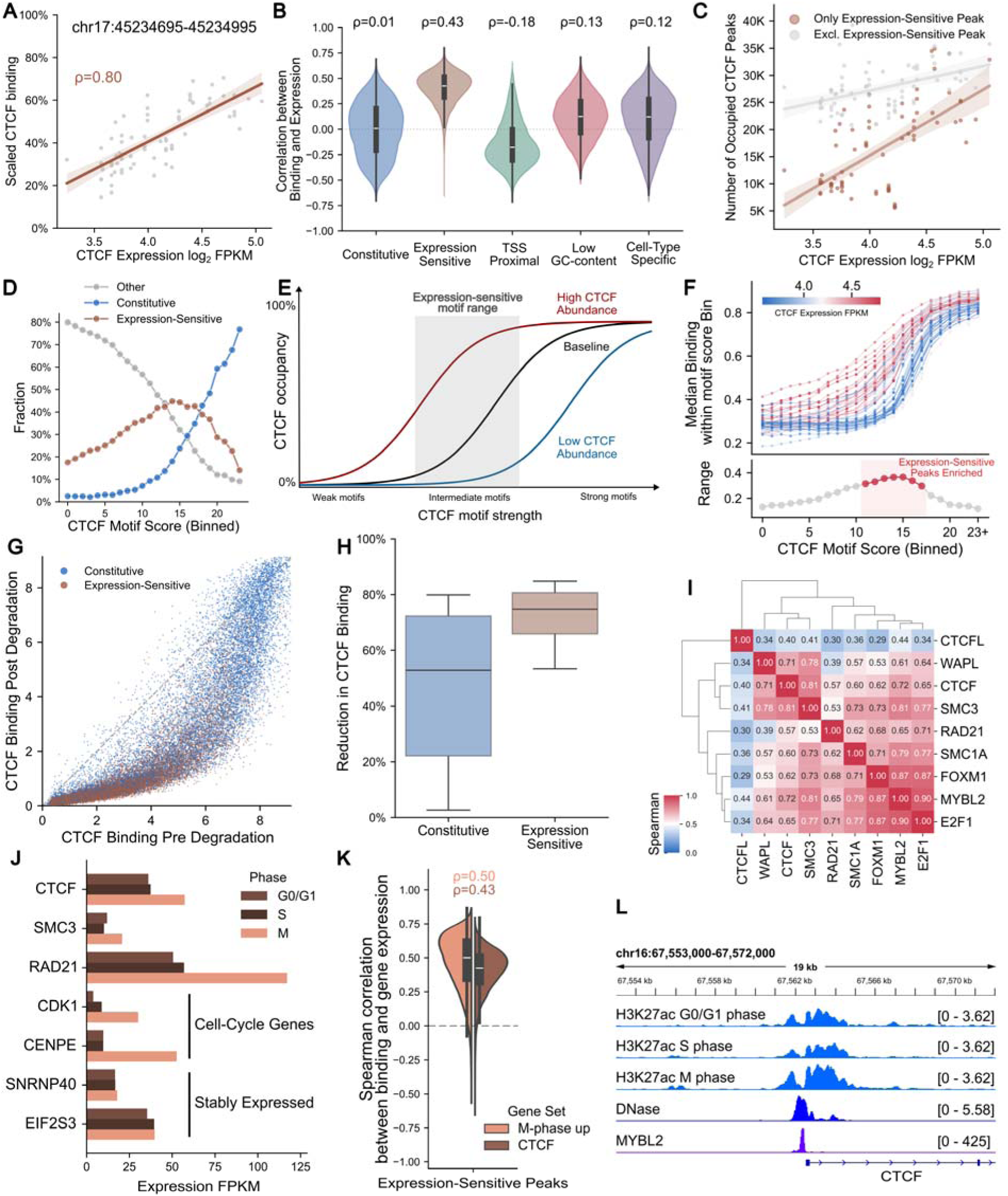
Identification of a large class of CTCF expression-sensitive binding sites with intermediate motif scores. (**A**) An example of a CTCF peak showing a strong Spearman correlation between CTCF binding and expression across datasets. (**B**) Distribution of binding-to-expression correlation across consensus peaks, stratified by peak categories. Almost all expression-sensitive peaks showed strong positive correlations. (**C**) Relationship between the number of occupied consensus peaks and CTCF expression level across datasets, grouped by whether peaks belonged to the expression-sensitive cluster. In samples with higher CTCF expression, more expression-sensitive peaks became occupied. In contrast, the number of peaks in other categories varied little. (**D**) Fraction of peaks, binned by motif scores, belonging to constitutive, expression-sensitive, and other clusters. A large fraction of peaks with moderate motif scores fell into the expression-sensitive cluster. (**E**) Schematic explaining why peaks with moderate motif scores tend to be most sensitive to changes in protein expression level. (**F**) Median CTCF binding strength across motif score bins in each ChIP-seq dataset. Each curve represented one dataset and was colored by CTCF expression in the sample. Bins with moderate motif score showed the largest occupancy variability across datasets. (**G**) CTCF occupancy at the constitutive peaks and the expression-sensitive peaks in the HUDEP-2 cell line, with and without CTCF auxin-induced degradation. (**H**) The fraction of reduction in CTCF occupancy after degradation in the HUDEP-2 cell line. Expression-sensitive peaks showed greater binding reduction than constitutive peaks. (**I**) Correlation among the expression levels of CTCF, cohesin subunits, and cell-cycle-related genes across biological samples. (**J**) Cell-cycle phased expression levels of CTCF and cohesin subunits. CTCF, SMC3, and RAD21 are all upregulated during the M phase. (**K**) Correlation distribution of expression-sensitive peaks between CTCF occupancy and CTCF expression or average expression of genes upregulated in M phase. Occupancy showed a stronger positive Spearman correlation with M-phase gene expression. (**L**) Phased H3K27ac binding and MYBL2 binding at the accessible CTCF promoter. The CTCF promoter has stronger H3K27ac modification and is bound by MYBL2.

Because regions with higher signals are more likely to be called peaks, we next investigated whether expression differences induced any differential peak calling in these clusters (**Extended Data Fig. 2B**). As expected, in the expression-sensitive cluster, samples with higher CTCF levels had more peaks called (ρ=0.56), whereas constitutive peaks were almost always bound, regardless of expression levels. Meanwhile, peaks in the cell-type-specific clusters were called primarily only in samples where those clusters were enriched. For instance, ESC-specific peaks were called only in embryonic stem cell lines. Overall, expression differences affected the number of peaks called, ranging from 30K to 70K, and total peaks called were positively associated with CTCF expression (ρ=0.55; **Extended Data Fig. 2C**). In addition, the expression-sensitive cluster is the main contributor to differences in the total peak count (**Fig. 3C**). The number of peaks assigned to all other clusters varied modestly, around 25K.

#### Expression sensitivity occurs at moderate-strength CTCF binding sites

We next investigated the properties of CTCF peaks more sensitive to expression. Across motif-score bins, the expression-sensitive cluster was relatively depleted in the high- or low-motif-score range but enriched in the intermediate range (**Fig. 3D**). Among peaks with CTCF motif scores between 11 and 17, more than 50% of these had Spearman correlations of 0.3 or higher with expression. Yet this number dropped to 10% among peaks in the highest motif-score bin (**Extended Data Fig. 2D**). This observation that expression-sensitive peaks tend to have moderate-strength sites is consistent with kinetic binding equilibrium (**Methods**). Fractional CTCF occupancy of a site depends on both the CTCF-DNA binding affinity (determined by motif score) and CTCF protein levels (determined by CTCF expression). At a fixed CTCF expression level, occupancy follows a sigmoid relationship with motif score, and as CTCF expression increases or decreases, the overall occupancy curve shifts left or right (**Fig. 3E**).

Thus, regardless of CTCF levels, peaks with high-affinity sites remained mostly saturated, and peaks with low-affinity sites stayed mostly unbound. In contrast, peaks with intermediate-affinity sites are quite sensitive to CTCF expression differences, with higher occupancy in samples where CTCF is upregulated. Although this modeling depends on nuclear CTCF protein concentration, we use gene expression as a surrogate. Median occupancy across peaks showed a sigmoidal relationship with motif scores (**Fig. 3F**), consistent with binding equilibrium modeling. Samples with higher CTCF expression levels have more left-shifted curves than those with low CTCF expression. Among motif score bins, the largest occupancy range across samples occurred in the moderate score range, bins with motif scores from 12 to 17, where the expression-sensitive cluster is enriched.

As an additional test of this hypothesis, we used the CTCF auxin-inducible degron (CTCF-AID) system to directly assess the expression sensitivity of these peaks.^6,35–38^ CTCF degron technology rapidly reduces CTCF protein abundance, allowing us to directly observe the effect of CTCF protein levels on binding strength within one cell type. In data from CTCF-AID in two erythroid cell lines, HUDEP2 and HEL,^35^ degradation induced a dramatic downregulation of CTCF binding at all sites (**Fig. 3G, Extended Data Fig. 2E**). Although all peaks showed weaker binding, the degree differed substantially between peaks in our constitutive and expression-sensitive clusters. The median binding reduction at constitutive peaks was 50% in HUDEP2, while most expression-sensitive peaks exhibited more than 70% reduction (**Fig. 3H**). In HEL, while peaks systematically lost more CTCF binding than in HUDEP2, the constitutive peaks were still more resistant to CTCF degradation (**Extended Data Fig. 2F**). The pronounced reduction at the expression-sensitive peaks further demonstrates that binding at these peaks is more reliant on CTCF expression levels, and that constitutive peaks, which are also stronger cohesin anchors, withstand CTCF depletion better.^36^

#### Differences in CTCF expression across samples is primarily due to different rates of proliferation

Finally, we investigated why expression for CTCF, an essential architectural protein,^39,40^ differs across samples, ranging from 8.8 to 33 FPKM. First, we noted that across ENCODE samples, CTCF expression was highly correlated with the expression of cohesin subunits SMC3 (ρ=0.81), RAD21 (ρ=0.57), and SMC1A (ρ=0.60), supporting their co-expression due to their shared role in genome organization (**Fig. 3I**). Further, CTCF expression also co-varies with WAPL (ρ=0.71), the cohesin unloading factor. This co-expression suggests that these proteins were likely regulated by the same biological process.

High CTCF expression was mostly observed in embryonic stem cells (H9=33 FPKM) and cancer cell lines (OCI-LY7=29), whereas fibroblasts had the lowest CTCF expression (IMR-90=8.8). Because the former are highly proliferative, we examined whether differential expression across the cell cycle might account for different bulk expression levels. Indeed, CTCF and cohesin subunits show higher expression in M-phase (**Fig. 3J**),^41^ similar to two known cell-cycle-regulated genes, CDK1 and CENPE. In particular, CTCF had 50% higher expression in M phase than in G0, and RAD21 had a 1-fold increase in M phase. Surprisingly, occupancy at the expression-sensitive peaks correlated better (ρ=0.52 vs. ρ=0.43) with the average expression of all genes upregulated in M phase (**Methods, Fig. 3K**), since averaging expression across many genes likely reduced noise and thus improved this correlation.

We also observed increased H3K27ac binding at the CTCF promoter in M phase, implying more promoter activity. This could be attributed to MYBL2 binding at the CTCF promoter (**Fig. 3L**), which is known to regulate many G2/M phase genes by binding directly at their promoters, such as CCNB1 and CDK1.^42^ Consistent with the postulated regulation, MYBL2 expression strongly correlates with CTCF expression (ρ=0.72) across ENCODE samples. Further, knockdown of MYBL2 effectively reduced CTCF expression by 25%.^43^

**Extended Data Figure 2:**
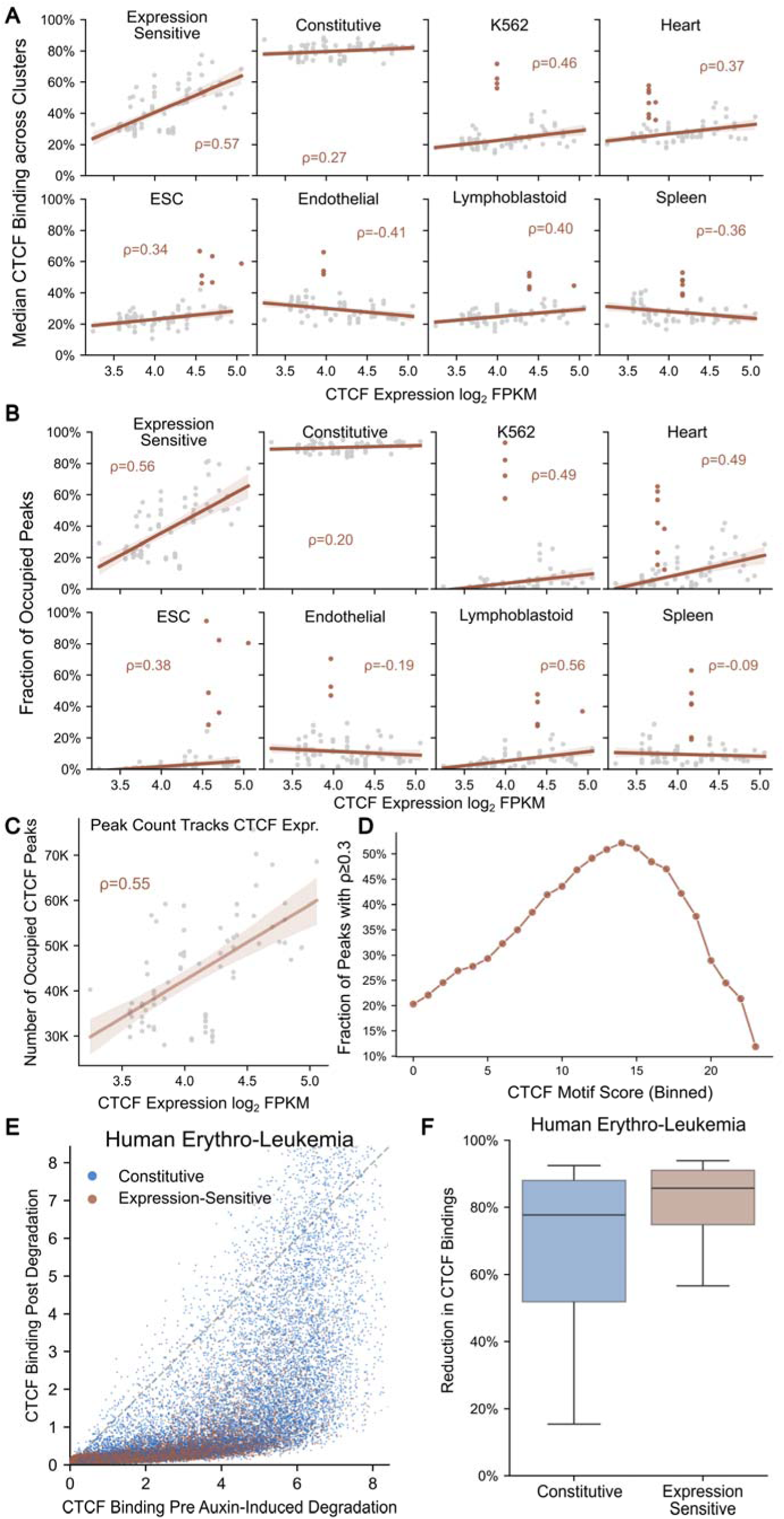
CTCF occupancy correlated most strongly with CTCF expression at the expression-sensitive peaks. (**A**) Across datasets, the average peak binding within each cluster was plotted against CTCF expression in that sample. Each dot represented one dataset. The expression-sensitive cluster showed a strong positive correlation between average binding and CTCF expression. In cell-type-specific clusters, binding correlated mildly with expression; the largest variation was observed in samples, which were highlighted in red, where these clusters were enriched. (**B**) Across datasets, the fraction of peaks occupied within each cluster was plotted against CTCF expression. The fraction of CTCF-occupied, expression-sensitive peaks correlated with CTCF expression. In cell-type-specific clusters, peaks were mostly occupied in samples where they were specifically enriched. (**C**) Across datasets, the number of CTCF-occupied peaks was correlated with CTCF expression levels. (**D**) Fraction of peaks, binned by motif scores, with Spearman correlations between CTCF binding and expression above 0.4. The fraction of such peaks peaked in bins with motif scores around 15. (**E**) CTCF occupancy at the constitutive peaks and the expression-sensitive peaks in the HEL cell line, with and without CTCF auxin-induced degradation. (**F**) The fraction of reduction in CTCF occupancy after degradation in the HEL cell line. Expression-sensitive peaks showed a greater reduction in binding than constitutive peaks.

### DNA methylation represses a subset of TSS-proximal CTCF peaks

Other clusters stood out by their genomic characteristics, because the majority of their peaks had high-GC content (>65%) and were proximal (<2 kb) to nearby TSSs, whereas other clusters had mostly distal peaks and a median of 50% GC content (**Fig. 4A**). We labeled these clusters as TSS-proximal C1, C2, and C3, which differed by other characteristics upon closer inspection. First, genes near C1 and C2 peaks were highly expressed with low variability across samples; however, C3-proximal genes were more variably expressed (**Extended Data Fig. 3A**). Gene ontology indicated that C1 genes were enriched in housekeeping programs, such as DNA replication and tRNA processing, yet C2 genes were more involved in cell-cycle-related processes (**Extended Data Fig. 3B**). In contrast, C3 peaks were proximal to many developmentally expressed genes with bivalently marked promoters.^38,44,45^ Finally, while all three clusters had well-above-genomic-background CpG content (1%), regions immediately flanking C3 peaks’ CTCF motifs were considerably more CpG-enriched (10%) than those next to C1 and C2 peaks (**Fig. 4B**).

**Figure 4:**
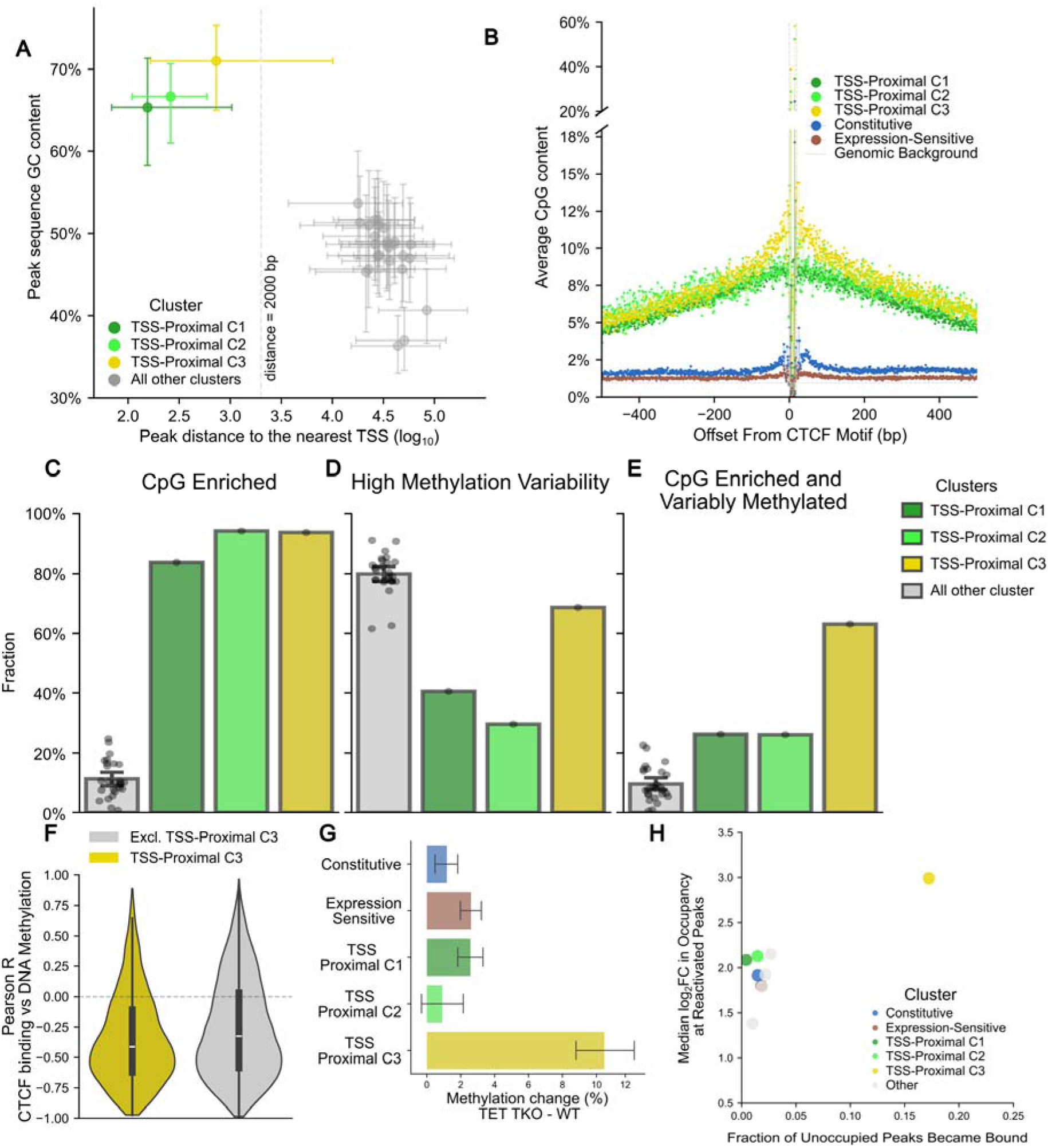
One set of TSS-proximal CTCF peaks was regulated by DNA methylation. (**A**) Distribution of peak clusters’ GC content and distance to the nearest TSS. Each dot denoted one cluster, with its position indicating the median GC content and median distance to the TSS. Error bars represented the 25th and 75th percentiles. Most peaks in the TSS-proximal clusters had GC content above 65% and fell within 2kb of the closest TSS. (**B**) Average CpG content within each CTCF peak cluster. Peaks were centered on CTCF motifs. CpG content flanking CTCF motifs in TSS-proximal clusters was significantly enriched compared with the genomic background or the CpG content flanking the peak motifs of constitutive or expression-sensitive clusters. Two CpG loci within the CTCF motif were also much higher than the background. (**C**) Fraction of peaks with more than 10 CG dinucleotides within each cluster. Each dot represented one peak cluster. (**D**) Fraction of peaks exhibiting variable methylation within each cluster. (**E**) Fraction of peaks with a large number of CG dinucleotides that are variably methylated within each cluster. (**F**) Distribution of Pearson correlation between CTCF occupancy and DNA methylation level across datasets for peaks satisfying both criteria. Most peaks showed a negative correlation, suggesting that DNA methylation antagonized CTCF binding. The median correlation in the TSS-proximal C3 group was significantly lower than in the other peaks. (**G**) Bar plot of the average change in DNA methylation between DNA demethylase triplet knockout and wild-type across peaks. CTCF peaks in the TSS-proximal C3 showed a large increase in methylation level. (**H**) Scatterplot between the fraction of peaks reactivated and the median log_2_ fold change in CTCF occupancy between DNA methyltransferase 1 knockdown and wild-type at the reactivated peaks. Compared with other clusters, occupancy at TSS-proximal C3 peaks increased more.

The high CpG enrichment in C3 peaks makes them ideal candidates for DNA-methylation-mediated inhibition of CTCF binding,^9–11^ Two positions within CTCF binding sites can contain CpG dinucleotides (across C3 peaks, at these two positions, 40% and 60% of CTCF motifs contain the CpG, respectively). Across all clusters, only TSS-proximal clusters had strong CpG enrichment (**Fig. 4C**). However, C1 and C2 peaks had limited methylation variability (**Methods**),^46^ likely because their neighboring genes were essential to all cell states and cannot tolerate repression of insulation (**Fig. 4D**, **Extended Data Fig. 3C**). In contrast, C3 peaks showed significantly higher methylation in cancer cell lines than in healthy tissue samples, and altogether only C3 had a majority of its peaks satisfying both criteria (**Fig. 4E**). In agreement with previous observations,^8,11,12^ among peaks with CpG enrichment and variable methylation, CTCF occupancy negatively correlated with methylation levels across samples (**Fig. 4F**), though, C3 peaks had modestly more negative correlations than others.

We next use genetic perturbation of the methylation modification machinery to show that C3 is unique among our clusters in its sensitivity to methylation. First, knockout of all three of DNA demethylases TET1, TET2, and TET3^47^ increased average methylation of C3 more than all other clusters (**Fig. 4G**). In addition, knockdown of DNA methyltransferase 1 (DNMT1)^48^ increased binding of previously unoccupied cell-specific C3 peaks, while also strengthening binding more at C3 peaks (**Fig. 4H**). These observations show that methylation-induced repression affects C3 more than other clusters.

However, methylation did not appear to act specifically on CTCF peaks, but on the broader region including promoters and TSSs. When we stratified TSS-proximal peaks by their relative position to nearby promoters, CpG content was consistently higher toward promoters (**Methods**, **Extended Data Fig. 3D**). When CTCF motifs were upstream of promoters, CpG level dropped sharply on the opposite side of promoters, suggesting CTCF bordered CpG-enriched regions. When we further stratified these peaks by the relative order of CTCF peaks, promoters, and TSSs, CTCF peaks consistently had the lowest CpG among the three (**Extended Data Fig. 3E**). Interestingly, promoters were more likely to be upstream of TSSs, and CTCF peaks were less likely to fall between promoters and TSSs. Overall, this result suggests that the variable occupancy observed at TSS-proximal CTCF sites likely reflects repression of the neighboring genes. In summary, while we find clear evidence that methylation of CpG interferes with CTCF binding, C3 explains a minority of CTCF variable occupancy sites, and therefore is not the dominant source of CTCF variability across cell/tissues.

**Extended Data Figure 3:**
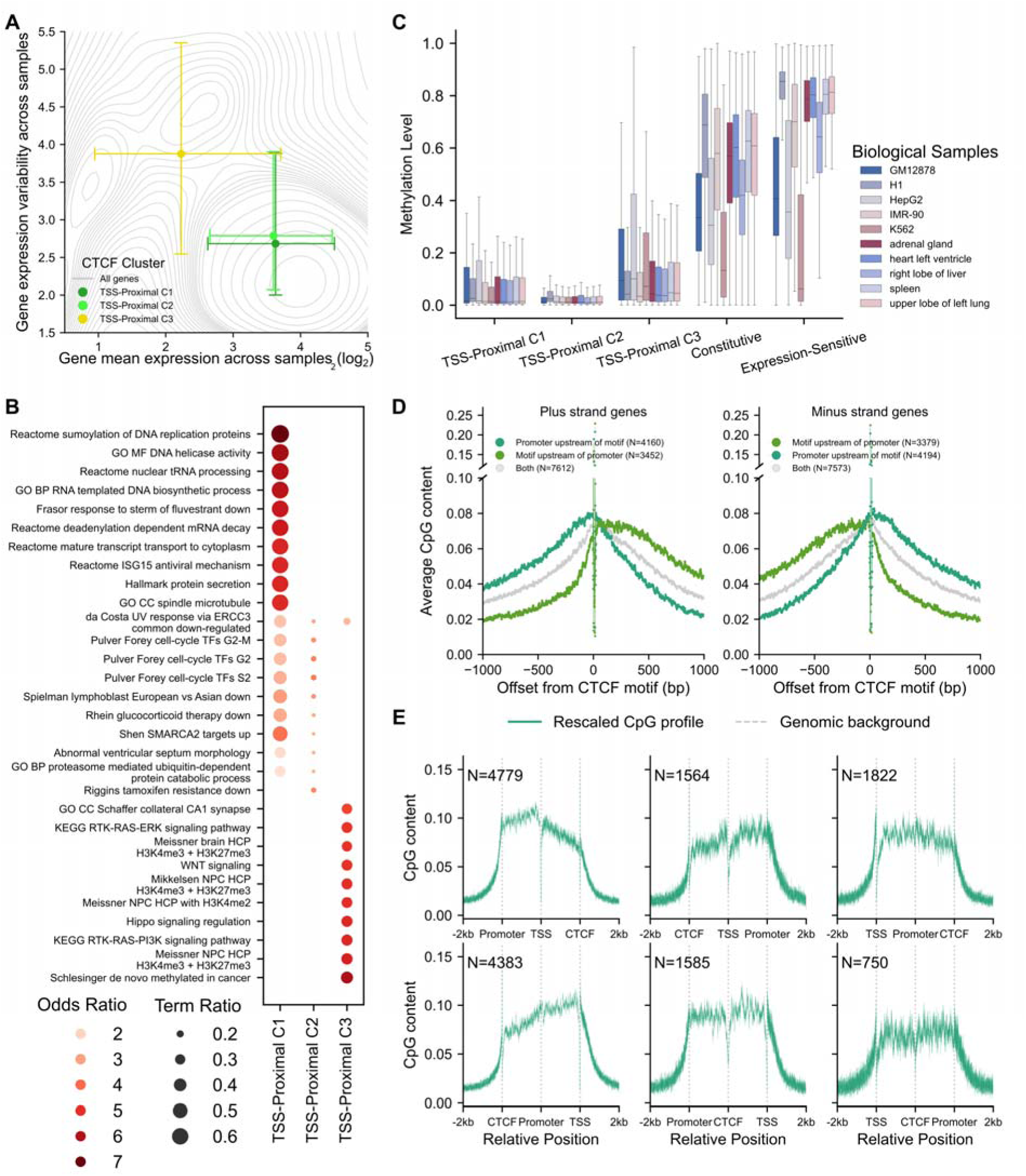
Characterization of TSS-proximal CTCF’s neighboring genes and CpG dinucleotide distribution. (**A**) Distribution of the mean and dispersion of gene expression for genes neighboring each TSS-proximal cluster. Each dot denoted one cluster, with its position indicating the median average expression and dispersion. Error bars represented the 25th and 75th percentiles. Genes adjacent to the TSS-proximal C3 peaks had lower average expression and were more variable across samples. (**B**) Gene ontology for genes that neighbored each TSS-proximal cluster. TSS-proximal C1 genes were enriched for the housekeeping terms, and TSS-proximal C3 genes were enriched for having bivalent promoters. (**C**) Distribution of methylation level across samples for peaks in the TSS-proximal, constitutive, and expression-sensitive clusters. Lower variability was observed among peaks in the TSS-proximal C1 and C2. (**D**) Across CTCF peaks proximal to a TSS with a promoter, average CpG content was computed for regions ±2 kb of CTCF motifs, stratified by relative position of the CTCF motif and its neighboring promoter. CpG content was enriched on the promoter side. (**E**) Across peaks proximal to an annotated TSS with a proximal promoter, we computed average CpG content in the regions ±2 kb of the CTCF motifs. The average CpG content was stratified by the relative position among the CTCF motifs, TSSs, and promoters. The averages were scaled such that the three elements were aligned across peaks within each of the six combinations.

### Cooperative binding with lineage-specific TFs modulates CTCF occupancy at cell-type-specific CTCF peaks

The final heterogeneous category of CTCF binding sites is cell-type-specific, with strong occupancy in a small subset of cell/tissues. We used our sequence-based ML modeling approach to identify TFs underlying these cell-type-specific CTCF peaks (**Fig. 5A**). We first trained gkm-SVM^20–22^ to learn predictive k-mer features that were enriched among the positive and negative sequences. We then applied gkmPWM,^49^ a motif-extraction method, to uncover TF motifs responsible for the observed k-mer enrichment or depletion (**Methods**). We chose gkm-SVM over currently popular deep neural networks because many clusters had too few peaks (around hundreds) for deep networks. We also didn’t opt for a multiclass model because we wanted to most sensitively determine TFs that discriminated classes. Therefore, we trained gkm-SVM in a pairwise, cluster-vs-cluster fashion and obtained a median cross-validated auROC of 0.87 (**Fig. 5C**). Cancer cell-line-enriched clusters were omitted because peaks that showed cancer-specific enrichment (e.g., K562-specific) often had copy number amplification (**Fig. 5B**). This amplification obfuscates peaks that were truly more occupied by CTCF from those potentially inflated by sequence read mapping.

**Figure 5:**
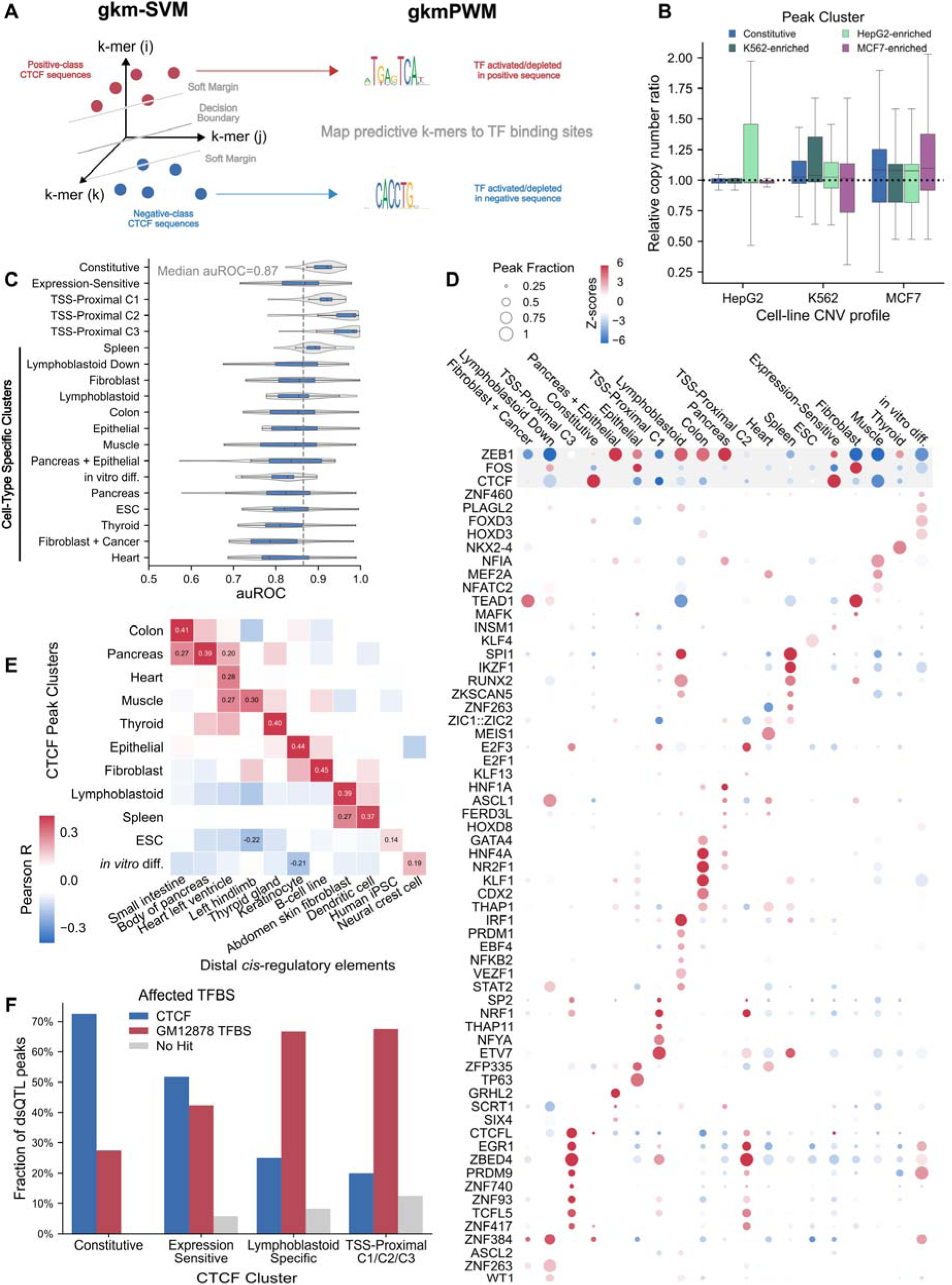
Cell-type-specific CTCF peaks had TF binding sites similar to those of distal enhancers of those cell-types. (**A**) Schematic of the identification of enriched and depleted TF binding sites with the gkm-SVM and gkmPWM pipeline. gkm-SVM embedded DNA sequences from two CTCF clusters into the gapped k-mer space and learned a sequence-space decision boundary that best separated the two sets. gkmPWM interpreted the trained gkm-SVM models by embedding TF motifs into the same gapped k-mer space and identifying the minimal, deduplicated set of motifs that collectively best support the learned decision boundary. (**B**) Distribution of relative copy number for peaks in the constitutive clusters and 3 cancer cell lines. A relative copy ratio of 1 means the cancer genome has 2 copies of the peak. Cancer-specific peaks were often found in amplified regions in that cell line (e.g., HepG2-specific peaks often had more amplified copies than constitutive peaks in HepG2). (**C**) Distribution of gkm-SVM’s auROC among pairs of CTCF clusters. auROC reported were derived from a five-fold cross-validation split along the chromosomes. (**D**) Dot plot showing motif enrichment and depletion for each CTCF cluster extracted by gkmPWM from gkm-SVM models. Dot color represented the strength of enrichment or depletion, and dot size indicated the fraction of peaks containing the corresponding motifs. Motifs below the gray box were ordered by tissue and cell type. (**E**) Correlation between gapped k-mer weights derived from the cell-type-specific CTCF peaks and those derived from distal *cis*-regulatory elements. CTCF peaks shared similar TF binding sites with enhancers from similar cell or tissue types. (**F**) Fraction of dsQTLs that altered CTCF binding sites and those that altered GM12878-specific TFs such as NFκB and EBF. dsQTLs in constitutive peaks mostly changed CTCF sites, and those in lymphoblastoid-specific peaks mostly changed non-CTCF binding sites.

From these clusters, gkmPWM attributed the predictive k-mers to motifs (**Fig. 5D**, **Extended Data Fig. 4A**) mapped to lineage-determining TFs, which we have shown are important for cell-type-specific enhancer activity.^50–60^ For instance, heart-related peaks were enriched for MEF2 motifs, and thyroid-specific peaks were enriched for Nkx motifs. Meanwhile, TEAD was predicted to be enriched in fibroblast-specific peaks^61^ yet depleted in epithelial-specific peaks.^51^ As expected, constitutive and expression-sensitive peaks had only CTCF motifs, with the former showing a stronger enrichment, and TSS-proximal peaks showed an enrichment for CTCFL.^14,62,63^ ZEB and AP-1 were enriched or depleted across many cell and tissue types, suggesting a possible interaction between CTCF and the TFs that recognize these two motifs. gkmPWM-derived Z-scores agreed with enrichment scores produced by motif-scanning tools (**Extended Data Fig. 4B**). Because distal enhancers are also cell-type-specific, we compared gkm weight vectors from models trained on these CTCF peaks to those from models trained on cell-specific distal enhancer elements. In terms of sequence composition, enhancers resembled CTCF peaks that were specific to the same cell type (**Fig. 5E**), implying that cell-type-specific TFs co-bound CTCF at these CTCF peaks.

These cell-type-specific peaks’ CTCF binding sites had much weaker affinities than those in constitutive or expression-sensitive peaks (**Fig. 2A**), so we hypothesized that the strong upregulation of binding at these sites was due to the presence of neighboring cell-type-specific TF binding. To answer this question, we examined lymphoblastoid dsQTLs, variants that induced changes in DNase I sensitivity.^64,65^ 33% of the 574 dsQTLs overlapped CTCF peaks; of the overlapping variants, 50% disrupted or created a CTCF motif (**Methods**). While 70% of dsQTLs in constitutive peaks altered a CTCF motif, only 20% of dsQTLs in cell-type-specific peaks altered CTCF motifs (**Fig. 5F**). dsQTLs contained within lymphoblastoid-specific peaks mostly changed GM12878-specific TF binding sites, such as PU.1 or RUNX3. These observations lend further support to the hypothesis that cell-type-specific peaks are bound by CTCF through interactions with cell-type-specific TFs.

**Extended Data Figure 4:**
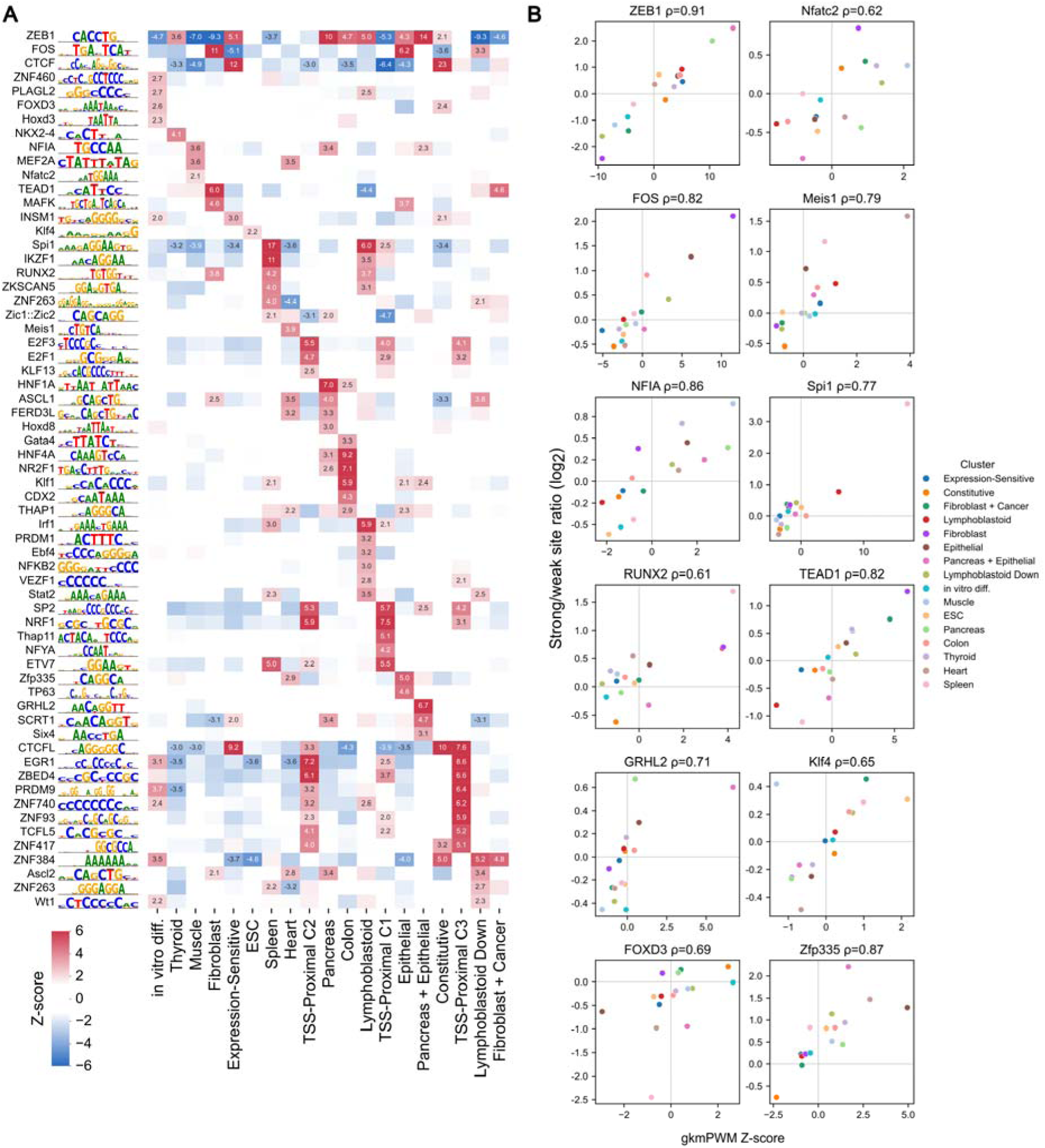
gkmPWM extracted enriched and depleted TF motifs. (**A**) Heatmap showing TF motif enrichment and depletion for each CTCF cluster extracted by gkmPWM. Colors represent the strength of enrichment or depletion, and scores for significantly regulated TFs in each cluster were annotated. Motif logos were displayed accompanying representative TF names. (**B**) Scatterplot showing correlation between motif enrichment predicted by gkmPWM and those derived from motif scanning. Motif enrichment was computed as the ratio of high-scoring motifs to low-scoring motifs. Many cell-type-specific TFs showed strong positive correlations between the two scores, offering an intuitive interpretation for gkmPWM motif scores.

### CTCF binding site affinity unifies distinct mechanisms regulating CTCF occupancy

In total, we described five categories of CTCF peaks, each regulated by a distinct mechanism. Since weaker peaks are always more variable (lower signal-to-noise), we evaluated the sensitivity of these categories on the threshold for peak detection. When we raised the threshold to reduce the consensus list from 126K to 82K peaks, the peaks still fall into these five categories, supporting the robustness of our findings to peak threshold (**Methods**, **Extended Data Fig. 5**). The cell-type-specific category was most sensitive to peak threshold, and shrunk by 50%, however, sequence features remained predictive of cluster membership (**Extended Data Fig. 5Q**). Additionally, methylation continued to modulate CTCF binding at peaks that were CpG-enriched and variably methylated (**Extended Data Fig. 5P**).

The five categories of peaks have binding sites of different CTCF motif strength (**Fig. 6A**), with cell-type-specific and TSS-proximal peaks showing much weaker binding affinity than others. Interestingly, though TSS-proximal and methylation-sensitive (previously labeled as TSS-proximal C3) peaks were both embedded in promoter contexts, the latter had much stronger binding sites. This suggests that we may only be able to detect methylation-mediated repression at high-affinity sites.

**Figure 6:**
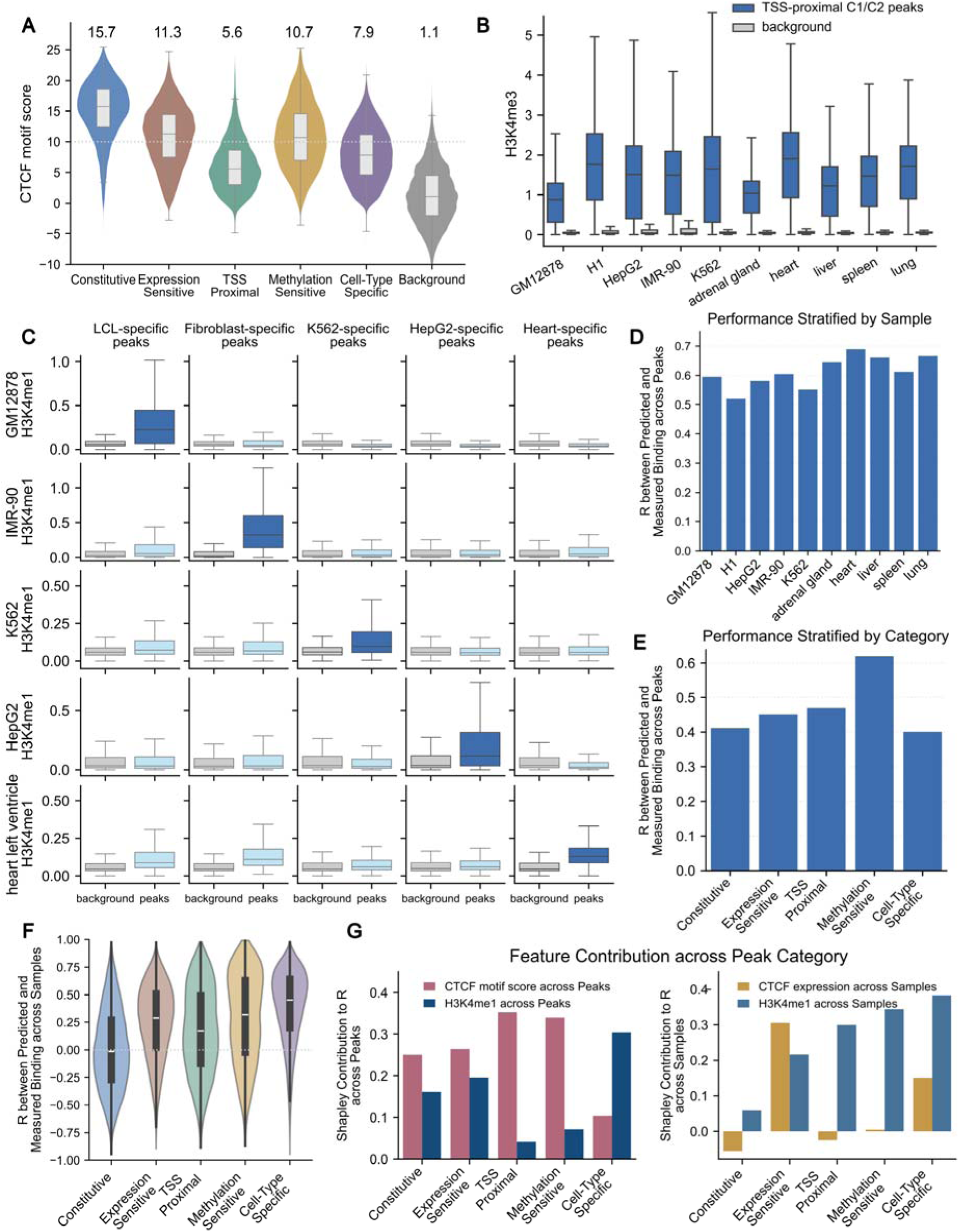
Peaks with low-affinity CTCF binding sites have strong histone marks, and chromatin features can predict CTCF occupancy on unseen samples. **(A)** Distribution of CTCF motif score strength across peak categories. The median motif score for each category was annotated above the violin plots. Background reflected randomly sampled 300bp regions. **(B)** Strength of H3K4me3 marks at TSS-proximal C1/C2 peaks versus that at GC-matched, CTCF-motif-score-matched genomic regions. The histone strength was computed as the average across the ±1 kb region. Across all samples, TSS-proximal CTCF peaks had higher H3K4me3 than matched background regions. **(C)** Strength of H3K4me1 marks at cell-type-specific peaks versus that at GC-matched, CTCF-motif-score-matched genomic regions. Cell-type-specific CTCF peaks had higher H3K4me1 only in samples where their binding was enriched. **(D)** Pearson correlation between measured and predicted CTCF occupancy across peaks, for 5 cell lines and 5 tissue samples. **(E)** Pearson correlation between measured and predicted occupancy across peaks, stratified by categories. **(F)** Pearson correlation between measured and predicted occupancy across 10 samples, for peaks in each category. **(G)** Chromatin features’ Shapley contribution to Pearson correlation across peaks and across samples, stratified by peak category.

Similar to how lineage-specific TFs co-bind with CTCF at cell-type-specific peaks, we hypothesized that promoter-associated TFs assisted CTCF binding at low-affinity TSS-proximal peaks. Compared with background regions having similar CTCF-DNA affinity, TSS-proximal peaks showed higher levels of promoter-marking histone modification H3K4me3 (**Fig. 6B**), supporting promoter-associated TF co-binding at these peaks. Meanwhile, cell-type-specific peaks had higher levels of enhancer-marking histone modification H3K4me1 relative to background, but only in samples with enriched binding (**Fig. 6C**). For instance, only HepG2-specific peaks displayed high H3K4me1 in HepG2.

In summary, CTCF constitutively occupies high-affinity sites via energetically favorable interactions. At moderate-affinity sites, elevated CTCF abundance can facilitate binding of otherwise unbound sites. As affinities decrease, binding becomes more sensitive to genomic contexts, specifically, cooperative binding assisted by cell-type-specific TFs or promoter-associated TFs. Therefore, at any given site, although occupancy was regulated by multiple mechanisms, intrinsic binding-site affinity determines the dominating mechanism modulating occupancy.

**Extended Data Figure 5:**
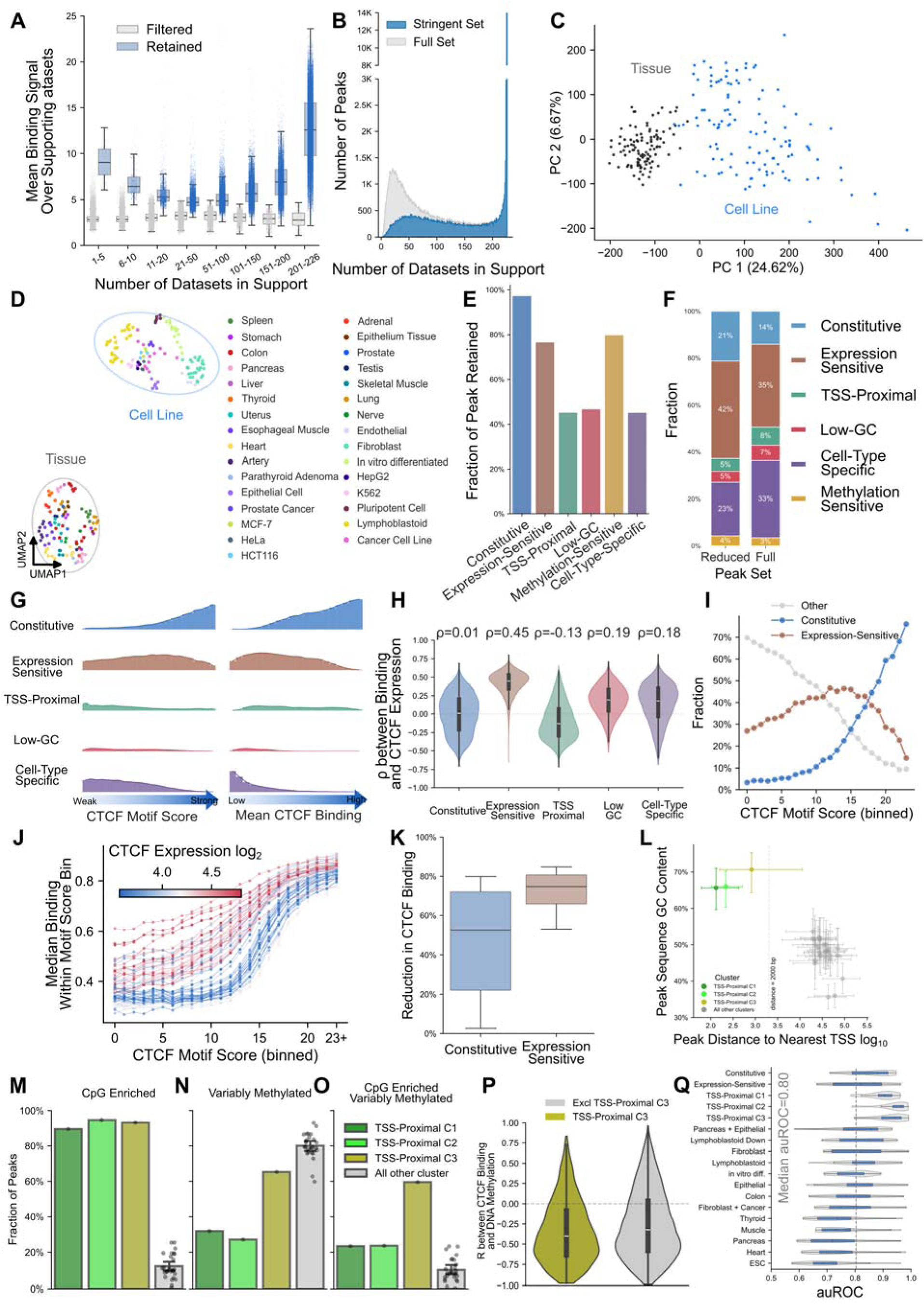
CTCF categories governed by different mechanisms reidentified on a smaller consensus peak list. (**A**) Peaks in the consensus list had higher average binding than the ones filtered out. After tightening the consensus list’s inclusion criteria, the differences in the binding strength distribution between the retained and removed sets increased. (**B**) Histogram of the number of datasets supporting each consensus CTCF peak. The number of variably bound peaks decreased. (**C**) PCA of CTCF ChIP-seq binding matrix on the smaller consensus peak list. Tissue-derived datasets remained separated from cell-line-derived datasets along the first principal component. (**D**) UMAP of CTCF ChIP-seq binding matrix on the smaller consensus peak list. Datasets assayed on similar biological samples remained clustered. (**E**) The fraction of peaks retained for each category. Almost all of the constitutively bound peaks remained. Most peaks in the expression-sensitive and methylation-sensitive categories were included, yet other categories had only half as many peaks as in the larger consensus list. (**F**) Fraction of peaks belonging to each category in the large and the small consensus peak list. (**G**) Fraction of peaks belonging to each category, binned by either CTCF motif scores or average CTCF binding strength. Constitutive peaks continued to occupy most of the highest-scoring bins and highest-binding bins. (**H**) Distribution of Spearman correlation between CTCF binding and CTCF expression level, stratified by peak categories. Expression-sensitive peaks still mostly showed strong positive correlation. (**I**) Distribution of consensus peaks among consensus, expression-sensitive, and other categories. Expression-sensitive peaks were enriched in bins with moderate-affinity binding sites. (**J**) Median CTCF occupancy binned by peaks’ CTCF motif score, for each ChIP-seq dataset. Compared with datasets with lower CTCF expression, those with higher CTCF expression tended to have stronger occupancy in a given motif-score bin. (**K**) The distribution of the percent reduction in CTCF occupancy post auxin-induced degradation in the HUDEP-2 cell line. Expression-sensitive peaks showed a larger reduction than constitutive peaks. (**L**) Median GC content and median distance to the nearest TSS for peaks in each cluster. TSS-proximal peaks continued to have significantly higher GC content than peaks in other clusters. (**M**) The fraction of 300bp peaks with 10+ CpG dinucleotides in each cluster. (**N**) The fraction of peaks experiencing variable methylation, measured by WGBS, across datasets. (**O**) The fraction of peaks satisfying both criteria with 10+ CpG dinucleotides and showing variable methylation. (**P**) Peaks satisfying both criteria had a negative correlation between DNA methylation and CTCF occupancy. (**Q**) The distribution of gkm-SVM auROC among CTCF clusters.

As an additional test to assess whether we have identified most of the mechanisms regulating CTCF binding, we used seven features to predict CTCF occupancy across consensus peaks in unseen samples (**Methods**). An XGBoost model was cross-validated by training on nine samples and evaluating on one held-out sample.^66^ This regressor achieved high correlation (R=0.5-0.7) between measured and predicted occupancy across peaks, in five cell lines and five tissue samples (**Fig. 6D**). When stratified by peak categories, the regressor remained accurate (R=0.4-0.6, **Fig. 6E**). Although our model was not trained on binding changes across samples, it accurately predicted binding variation on some peaks, with the exception of constitutive peaks where variation largely reflects noise. Cell-type-specific peaks exhibited a median correlation of 0.5 (**Fig. 6F**). Shapley feature contributions indicated that CTCF motif score was the most important feature for predicting occupancy, consistent with the strong correlation observed between the two (**Methods**, **Fig. 6G**). H3K4me1 overtook motif score and was the strongest predictor for cell-type-specific peaks. Meanwhile, H3K4me1 levels were influential for predicting binding changes across samples, and CTCF expression levels best modeled binding variation at expression-sensitive peaks.

### Differential occupancy predicts differential contact frequency and expression

The most likely impact of variable CTCF binding is that it can contribute to changes in enhancer-promoter contact frequency and gene expression through modulation of TAD structure. Having established distinct mechanisms modulating CTCF occupancy, we next show that this variable CTCF binding affects CTCF-mediated chromatin loop strength. As a specific example, near BUB3, a gene involved in cell division, contact frequency exhibits strong correlation with occupancy (R=0.78) between two CTCF peaks (**Methods**, **Fig. 7A**). Across 33 samples, increased binding strength at the convergent pair is accompanied by 10-fold increase in Hi-C-measured^2,67^ contact frequency. The emergence of these peaks also allowed the constitutive peak at 123.15 Mb to form a loop, nested within the larger one. Loop strength displayed a moderate correlation with BUB3 expression (R=0.44).

**Figure 7:**
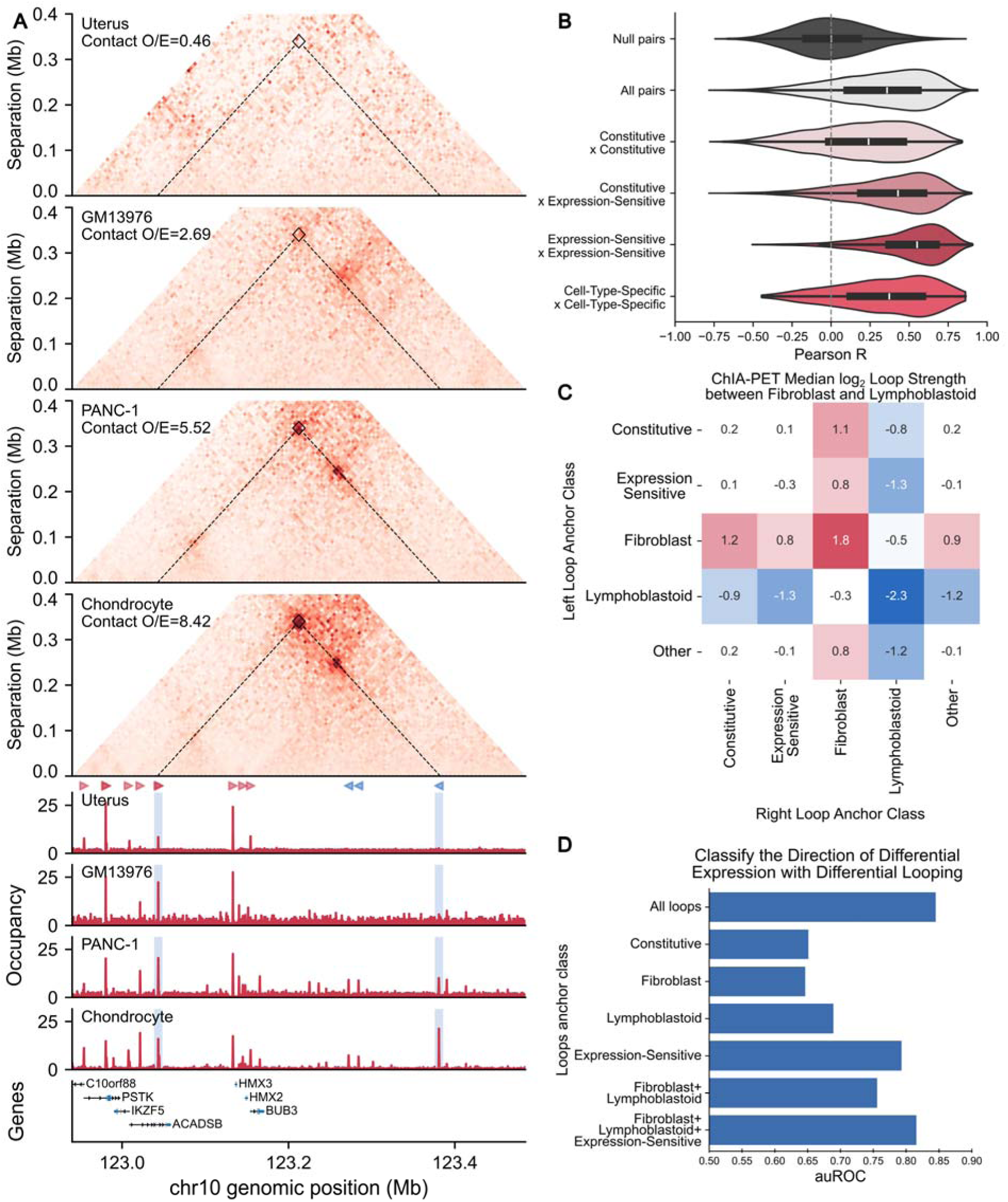
Differential CTCF occupancy affected differential contact frequency, which predicted differential expression. (**A**) An example gene locus on chromosome 10 where, across samples, CTCF occupancy covaried with contact frequency between the two CTCF peaks. Contact between the two peaks was boxed in squares on Hi-C contact maps, and the interacting CTCF peaks were shaded in blue on ChIP-seq tracks. Arrows indicated binding site orientation within each CTCF peak. (**B**) Distribution of correlation between CTCF occupancy and contact frequency across Hi-C data. “All pairs” described all convergent interacting pairs, and “Null pairs” described a size-matched set of non-interacting pairs. We highlighted some subsets of peak pairs stratified by anchor category, such as constitutive, expression-sensitive, or cell-type-specific. Occupancy was computed as the sum of both anchors’ CTCF binding. (**C**) Median log_2_ fold change in loop strength between fibroblasts and lymphoblastoid, stratified by loop anchor cluster. Colors reflected fold-change strength, with red indicating loops were stronger in fibroblasts and blue indicating loops were weaker in fibroblasts. (**D**) auROC of predicting the direction of genes’ differential expression from loop strength. Strength was defined as the sum of all loops enclosing a gene’s TSS. We used the top 1,000 expressed genes from fibroblasts and lymphoblastoid (2,000 total) for this classification task. We then subsetted loops by anchor cluster membership. For each loop subset, we recomputed log_2_ fold change in loop strength and predicted the direction of differential expression.

Across all peak pairs that form strong loop in at least one sample, we observed a moderate correlation (R=0.36) between occupancy and contact, whereas the null distribution showed a median R of −0.0001 (**Fig. 7B**). As expected, constitutive peak pairs have a weaker correlation of 0.24. In contrast, expression-sensitive peaks’ occupancy strongly influenced loop strength. The median correlation increased to 0.43 for loops formed between constitutive and expression-sensitive peaks, and to 0.55 for loops established between two expression-sensitive peaks. In addition, cell-type-specific peak pairs showed a moderate correlation (R=0.37). Thus, expression-sensitive CTCF peaks have the strongest effect on cell-type specific chromatin looping.

In addition to Hi-C, which measures interactions among all CREs, ChIA-PET directly quantifies loop strength between CTCF peaks.^68^ Between fibroblast and lymphoblastoid cell lines, loops anchored by two constitutive or two expression-sensitive peaks exhibited no variation (**Fig. 7C**). However, for each additional fibroblast-specific anchor, median loop strength increased twofold. In other words, loops where both anchors were in the fibroblast-specific cluster showed fourfold enrichment. Similarly, for each additional lymphoblastoid-specific anchor, median loop strength decreased 2-fold. Interestingly, among loops with one fibroblast-specific anchor and one lymphoblastoid-specific anchor, loop strength remained similar between samples.

Moreover, stronger chromatin loops are associated with gene upregulation (auROC=0.84). For a given gene, increased strength for loops enclosing the gene indicated likely upregulation in that cell type (**Methods**, **Fig. 7D**). In contrast, weakened loops are associated with downregulation in fibroblasts. Predictive power reduced substantially when we only considered constitutive peaks (auROC=0.65); however, using only lymphoblastoid-specific, fibroblast-specific, and expression-sensitive loops yielded similar accuracy (auROC=0.82). The significance of expression-sensitive loops likely reflected differences in CTCF expression between the two cell types.

## Discussion

In this study, we developed and demonstrated the utility of an analysis framework to identify novel mechanisms underlying variable CTCF occupancy. Leveraging diverse tissue and cell types, we partitioned peaks into groups with distinct binding enrichment patterns across datasets. Each pattern can be attributed to a biological mechanism. In total, we labeled 28 clusters in five broad categories and explained each distinct category. In particular, we identified CTCF expression as responsible for the largest group of variable-occupancy peaks, and we showed that CTCF expression differences are cell-cycle dependent and reflect differences in proliferation rates. For example, highly proliferative samples such as embryonic stem cells and cancer cell lines tend to have higher CTCF expression, occupancy at CTCF expression-sensitive peaks, and concomitant variable strength chromatin loops.

Outside of the constitutive cluster, whose peaks had high-affinity binding sites, other clusters exhibited variable occupancy and had peaks with lower-affinity sites. Because binding sites were weaker, these peaks were more susceptible to cell-type-specific influences, leading to variation in binding across tissue and cell lines. We showed that these sites were occupied either because of high CTCF expression or favorable genomic contexts set by nearby promoter- or enhancer-associated TF binding. The presence of TF binding sites was supported by sequence-based ML modeling. gkm-SVM models attained high predictive performance across cell-type-specific clusters. These models are easily interpretable through gkmPWM.^49^

CTCF binding-site strength determines the dominant mechanism of sites within each cluster. While we only analyzed CTCF here, the biophysical mechanisms we identified (protein abundance, methylation, and TF cooperative binding at flanking sites, apply to other TF binding) likely apply to other TF binding modulation as well. However, because CTCF has a much longer binding site than most other TFs do, we have reported that other TFs are more reliant on co-factors than CTCF and cohesin subunits,^21^ and their variability is thus likely much more dependent on co-factor-dependent changes in local accessibility than CTCF.

We demonstrated that our framework explained most of the variability in occupancy on held-out samples with just seven features. We did not include ATAC signal in this model because chromatin accessibility could be a result of, rather than a mechanism of binding.^69^ Finally, we observed that contact frequency between two CTCF peaks correlated strongly with occupancy at the peaks, and that these stronger chromatin loop structures are associated with upregulation of enclosed genes. In contrast to our modeling, previous computational methods that predict Hi-C contact from DNA sequence often fail to generalize to unseen cell types.^70^ This may partly be because sequences do not capture CTCF abundance, so models struggle to predict contacts anchored by expression-sensitive peaks, which make up close to 40% of the consensus peak list.

Consistent with previous reports, our findings found that DNA methylation antagonized CTCF binding, and that cell-type-specific binding enrichment mostly resulted from cooperative binding facilitated by cell-type-specific TF co-binding. However, we showed that methylation-mediated repression only strongly affects one small group of peaks that were largely TSS-proximal, accounting for less than 10% of all CTCF peaks. Nonetheless, DNA methylation remains a crucial mechanism regulating TF binding in embryogenesis.^12^ We attributed this refined estimate to the much larger ENCODE CTCF ChIP-seq datasets and our framework’s ability to jointly consider all putative mechanisms.

Currently, our framework does not attribute differential TF occupancy to mutations in *cis*-regulatory elements that are common in cancer cell lines or tumor samples. While we have observed that copy number variation contributes to some observation of what appears to be variable occupancy, large copy number variation can be factored out. However, our approach does not try to account for single-nucleotide variants, which could also contribute to altered TF binding or chromatin accessibility at a small number of peaks.

The identification of these mechanisms of variable occupancy should inform future modeling efforts. While we have applied our framework only to variable CTCF binding in this study, it could be easily extended to other TFs, such as REST, EZH2, and CEBPB, though its utility relies on a large number of datasets spanning different cell types or cell states. In addition, this framework could be applied to analyze ATAC-seq, for instance, to understand differential peak activation across a cohort of patients at various disease stages.

## Software and Data Availability

The scripts and data needed for reproducing figures presented in this manuscript is made available through Zenodo under the link: https://zenodo.org/records/24325166?preview=1&token=eyJhbGciOiJIUzUxMiJ9.eyJpZCI6IjM1Y2FiNGVhLWRmMzQtNDViMS04YjdhLTZhNDY0OTE2YTRlOCIsImRhdGEiOnt9LCJyYW5kb20iOiJiOTdiZTNmNjUwZjFhYWJhMWZhODhlOWMyMWNlMTkxYiJ9.54qgAfcIGud1DhS9thRFPj4eZjXMVLjEydF4ixyWy0Qx-glulaJoqu2jErvLUckTdw7cj3BXZ6tm6chJei-P6g

CTCF, SMC3, RAD21, JUN, JUNB, JUND ChIP-seq, RNA-seq, WGBS, H3K4me1 ChIP-seq, H3K4me3 ChIP-seq, H3K27me3 ChIP-seq, Hi-C, and ChIA-PET are available on the ENCODE portal (accessions provided in the supplementary tables). Cancer cell-line copy number variations are available on DepMap under 26Q1 release. Additional sequencing data are available at GEO, under accession No. GSE281734 (CTCF-AID), GSE94479 (cell-cycle phased RNA-seq and H3K27ac ChIP-seq), GSE217752 (CTCF ChIP-seq after DNMT1 interference), and GSE89728 (WGBS after TET triple knockout). TF motif PWMs are available on JASPAR.

## Acknowledgements

We thank Milad Razavi-Mohseni, Andrew Rojnuckarin, Dustin Shigaki, and Jin Woo Oh for their helpful discussion and valuable insights. In particular, we thank Milad for compiling ENCODE RNA-seq and WGBS data, Andrew for assistance with ChIA-PET analysis and clustering discussions, and Dustin for assisting with gkmPWM utilization and optimization. We also thank Benjamin Albert, Runxin (Wendy) Wu, Alejandro Chavez, and Danwei Huangfu for useful discussions and feedback on the manuscript. We especially appreciate Benjamin and Runxin for discussing many analyses. We acknowledge funding from the following: National Institutes of Health grant R01HG012367 (M.A.B.) and U01HG012051 (M.A.B.).

## AI Disclosure Statement

AI was used to improve some visualizations and to polish the manuscript grammar. Both authors have carefully reviewed and verified the accuracy of all text, all scripts for which there was AI input, and all figures.

## Author Contributions

M.A.B. conceptualized, supervised, acquired funding, and provided computing resources for this project. Y.Y. developed methodology, analyzed data, and produced figures. M.A.B. derived the kinetic binding equilibrium equation and constructed TSS-to-promoter mapping. Y.Y. wrote the manuscript. M.A.B. reviewed and edited the manuscript.

## Methods

### CTCF ChIP-seq data processing

All 302 CTCF ChIP-seq datasets were obtained from the ENCODE consortium (encodeproject.org).^71^ The complete list of dataset accessions is available in the supplement (**Supp. Table 3**). We downloaded the filtered alignment files in BAM format. Two datasets (ENCSR336PTS and ENCSR780OZE) had alignment files labeled as redacted alignments, rather than just alignments. These files were used as input to MACS2 (Model-based Analysis of ChIP-seq, v2.2.9.1).^72^ MACS2 `callpeak` was run on all replicates’ alignment files together, with default statistical cutoffs (-nomodel -B -SPMR), to generate peak calls. The command generated two signal tracks named “lambda_control” and “treatment”. The former stored background computed from pileup, and the latter stored reads. Only reads mapped to major chromosomes were retained. MACS2 `cmpbdg` with option (-FE) was used to convert raw read signal to fold enrichment over estimated background. Tracks were finally stored in bigwig format.^73^

### CTCF ChIP-seq quality control

Inspired by gkmQC,^19^ we similarly quality-controlled CTCF ChIP-seq with gkm-SVM.^20–22^ For each dataset, we first took the top 20,000 called by MACS2. Peaks were ranked by the number of reads at summits and then partitioned into 8 sets of 2,500 peaks each. The first set contained the strongest 2,500 peaks, the second contained the next 2,500 strongest peaks, and so on. Each peak set was matched with an equal-sized set of GC-matched, repeat-fraction-matched background sequences. Then, gkm-SVM models with default parameters were trained (https://github.com/Dongwon-Lee/lsgkm). We used five-fold cross-validation auPRC (area under the precision-recall curve) as the predictive performance metric, where sequences were split along chromosomes to prevent data leakage.^64^ Thus, each dataset ended up with 8 such auPRC scores, except for the ones where MACS2 called fewer than 20,000 peaks. For each of the 8 bins, we pooled auPRC from all 302 datasets and computed the lower bound of the interquartile range (IQR) as the cutoff. The lower bound was calculated as Q1 - 1.5 * (Q3 - Q1). Bins with stronger peaks had higher cutoffs. Datasets with all 8 auPRC above the corresponding cutoffs were considered high-quality, and the rest were filtered out.

Based on donor information stored in the experimental metadata, we further removed datasets that included multiple donors. After this step, we had 240 experiments. Finally, we removed experiments that showed low cross-replicate correlation. We computed this correlation as follows. For each dataset, we computed the average signal for all 1 kb windows, tiling along the entire genome. Signal was retrieved from the fold enrichment over background bigwig tracks. We then constructed a signal matrix with 3.088 million windows and 240 experiments. We took the top 30,000 windows, ranked by average signal across experiments. We then computed cross-experiment Pearson correlation on the 30,000-by-240 matrix. For biological samples with more than one dataset, we kept removing the most dissimilar dataset until the average correlation for that biological sample reached 0.8. For biological samples with only one dataset, we examined its correlation to other datasets that characterized similar samples. For instance, the heart left ventricle and heart right ventricle were both categorized under heart samples. We removed singleton datasets that did not reach a Pearson correlation of 0.8 with any other dataset characterizing a similar biological sample.

### Consensus peak list construction

From all high-quality experiments, summits (stored in .narrowPeak files) with p-values lower than 0.001 were extended into 300-bp peaks (150 bp upstream and downstream). Datasets had an average of 87K peaks, with a standard deviation of 23K. These peaks were then pooled together. Peaks overlapping by at least 149 bp were stitched into regions, yielding 2.6 million regions. Of these, 523K were supported by at least three experiments. For each well-supported region, the center was computed as a weighted average between original peak centers and fold-enrichment signals. The region centers were then extended to 300-bp windows. Windows that overlapped the ENCODE blacklist were removed, and STORM was used to scan for the strongest CTCF motif scores within these windows.^29^ CTCF position-weight matrix (PWM) was obtained from JASPAR under accession MA0139.1. STORM was run with the option (-m), which reports the binding site instance with the highest score. The software outputs the start position of the highest-affinity binding site, the binding site orientation, and the motif score. The motif score was subsequently used for binding peaks, and the orientation was subsequently used for CpG content analysis.

For each window, we subsequently obtained the average fold-enrichment signal from all high-quality experiments. We then applied a second round of signal-based filtering. Each peak was required to have a signal at least 25% of the 3000th-highest signal in at least 3 experiments. After filtering for high signal, above-0 motif score, and no blacklist overlap, we ended up with 126,579 peaks in our consensus list. From these consensus peaks, we recalculated peak centers as a weighted average of center and signal, but only from datasets with high enough signals (25% of the 3000th-highest). Upon calculating new peak centers, they were extended back to 300 bp again, and average signals were re-extracted based on the final window positions. The final binding matrix consisted of 126K peaks and 226 experiments.

### PCA and UMAP embedding

We first normalized the raw binding matrix (with dimensions 126,579 by 226). We computed the average binding signal across all ubiquitously bound peaks. These were the peaks with sufficiently high signals across all datasets. The matrix was normalized such that the average signal across these high-signal peaks became equal. The normalized binding matrix was then standardized so that all peaks had an average binding of 0 and a standard deviation of 1. Scikit-learn PCA was used to solve for the first three principal components.^74^ UMAP (v0.5.7) embedding was computed in the correlation space (n_neighbors=10 and min_dist=0.4).^75^ Clustering by sample remained largely robust across parameter choices. We chose this parameter combination to avoid visualizing overly tight clusters. Both PCA and UMAP low-dimensional representations were recomputed on a subset of datasets generated by the Bernstein lab using the antibody labeled as ENCAB000AXXX, sourced from Millipore.

### RNA-seq data processing

RNA-seq gene quantification was downloaded from the ENCODE portal. Expression values, in units of FPKM, for protein-coding genes across datasets were merged. After adding a pseudocount of 1, the upper-quartile normalization was performed so that the 75th percentile across datasets was equal. Afterward, to minimize technical batch effects, we only retained experiments that characterized a specific cellular component, received no treatment other than DMSO, were labeled as total RNA-seq, and were generated by the Wold lab. Intersecting these datasets with the biological samples for which we had CTCF ChIP-seq left us with 72 RNA-seq datasets, corresponding to 75 CTCF ChIP-seq datasets (**Supp. Table 4**). The matching process required that both the biosample term name and life-stage annotation matched. We then merged the RNA-seq datasets to reflect gene expression per sample type, such as spleen or adrenal gland. Datasets were merged by taking the median expression value for a gene. For many downstream analyses involving CTCF expression level, we assigned all ChIP-seq datasets characterizing a sample the same median CTCF expression value.

### Correlation analysis between CTCF occupancy and CTCF expression

RNA-seq processing procedure was described in a previous section. We calculated the Spearman correlation coefficient between CTCF occupancy and CTCF expression level on matched samples. CTCF ChIP-seq datasets characterizing the same biological sample were paired with the same CTCF expression level, which was the median expression value calculated from all RNA-seq data characterizing that sample. We then permuted the columns of the binding matrix and recalculated the Spearman correlation for the null distribution. For each peak’s binding-expression correlation, we then assigned a nominal p-value based on the permutation test. FDR was subsequently calculated, and we deemed a peak to be expression-sensitive if it had a positive Spearman correlation with FDR below 0.01.

We also calculated Spearman correlation between expression and occupancy across the entire cluster. For that, we first transformed the normalized binding signal to log_2_ scale, after adding a pseudocount of 1. Then, for each dataset, we scaled the log signals across peaks so the minimum and maximum values became 0 and 1. Then, for each cluster, we computed the median signal across all peaks in that cluster. Thus, for each cluster, we had one value representing binding strength for each dataset, and we correlated this median binding with that dataset’s CTCF expression.

### Peak-wise clustering

We transformed the normalized binding matrix to fold-change relative to the average signal. To do so, we divided the matrix by its row mean, adding a pseudocount of 1 to the numerator and denominator before division. Row means showed each peak’s average binding strength across all datasets, representing baseline occupancy. The fold-change-relative-to-baseline matrix was stored as an AnnData (v0.11.1), where we then used Scanpy (v.1.10.3) to compute nearest neighbors.^25^ The neighbor calculation was done on the whole matrix (not reduced principal component representation), across the correlation metric. The nearest 40 neighbors were identified for each peak. We then performed Leiden clustering,^24^ using the Constant Potts Model (partition_type=CPMVertexPartition), a non-directed graph, and a high resolution to promote somewhat comparable-sized small clusters. Resolution was set to 4 * 40 / 126,579. In other words, each clustered community must have 4-fold the baseline graph edge density (n_neighbors / number of peaks).

The strict edge-density requirement produced many clusters, which we then manually merged. We let the Leiden algorithm under-cluster rather than over-cluster. We used mostly cell-type-specific binding enrichment/depletion patterns to merge clusters, though features such as peak GC content, correlation to CTCF expression level, and peak normalized occupancy across datasets also helped. We combined clusters with more than 55% expression-sensitive peaks to form the final expression-sensitive cluster. Clusters smaller than 400 peaks were discarded, and we ended up with 28 clusters, representing 96% of the consensus peak list.

### SMC3 and RAD21 ChIP-seq data processing

SMC3 and RAD21 ChIP-seq datasets were obtained from the ENCODE consortium (encodeproject.org). As described previously for CTCF ChIP-seq, we used MACS2 for peak calling, signal track generation, and fold-enrichment track calculation. Then, for each dataset, for each peak, we extracted the average signal within the window. This produced a matrix consisting of 126K peaks by 10 experiments (**Supp. Table 5**). We then calculated normalization factors to make all datasets have the same average. Afterward, we extracted the per-base-pair signal for 2000-bp windows centered on each peak center for each dataset and normalized it with the precomputed normalization factor. Datasets characterizing the same biological samples were then averaged and visualized. Peaks within the tornado plot were sorted within each category, for each of the three samples. The tornado plot color bar was clipped to the 1st and 98th percentiles.

### JUN ChIP-seq data processing

JUN, JUNB, and JUND ChIP-seq datasets were obtained from the ENCODE consortium. As described previously for CTCF ChIP-seq, we used MACS2 for peak calling, signal track generation, and fold-enrichment track calculation. We skipped gkmQC-style quality control for these datasets but removed those with low cross-replicate correlation. No ChIP-seq dataset characterized more than one donor. As previously described, we then created a consensus peak set. We pooled all peaks with p-values < 0.001, stitched them into regions, and accepted regions with more than 3 supports, yielding 156K regions. We then scored them against the JUN position-weight matrix (JASPAR MA0099.4) using STORM. We considered these to be high signal if the average signal was at least 40% of the 3000th-highest signal in at least 3 experiments. Finally, we retained only peaks with high signal, an AP1 motif score > 0, and no overlap with the ENCODE blacklist. We then counted the number of peaks present in each dataset, by only considering those peaks with high signal to be occupied.

### Kinetic modeling of CTCF-DNA interaction

The equilibrium binding and dissociation of CTCF and DNA can be described by the dissociation constant:

Because we have a fixed amount of CTCF and DNA, we define two constants: [Total CTCF] = [CTCF] + [CTCF–DNA] and [Total DNA] = [DNA] + [CTCF–DNA]. Substituting into the dissociation constant definition, we get

Writing in terms of the fractional occupancy, θ:

Thus, we have

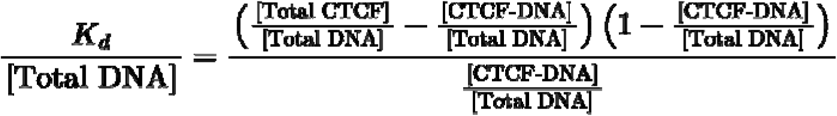

Now, we could solve for this equation, where if we simplify the notation with

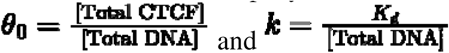

we then have:

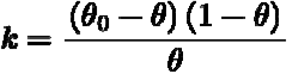

and therefore:

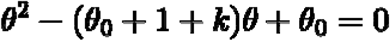

Solving the quadratic, we have

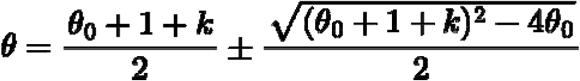

Where only the minus version is a valid solution

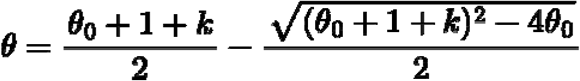

Now, because motif score is linearly related to −ln(K_d_) and −ln(k),^26,27^ fractional occupancy depends sigmoidally on motif score.

### CTCF auxin-inducible degron data processing

We downloaded the CTCF-AID study’s sequencing files from NCBI GEO and trimmed Illumina read adapters with cutadapt (v5.2).^76^ Reads from two replicates of the control and the treatment group (total of 4 sets of paired-end reads) were separately mapped to hg38 using Bowtie2 (v.2.5.4).^77^ We only retained the ones mapped to major chromosomes. Upon converting them to BAM format, we combined replicates for MACS2 peak calling and signal track generation. We then extracted average signals within windows defined by our consensus peak list. No additional normalization was applied beyond the library size normalization (--SPMR) in MACS2. Percent signal reduction was calculated as the signal difference between control and treatment over the control group signal. We then used BedTools `intersect` to subset our consensus peak list to peaks overlapping MACS2-called peaks in the control group, focusing only on peaks present in HEL and HUDEP2 (50K and 55K peaks, respectively). Then, for each cell line, we visualized the average signal at peaks in the control versus treatment group. We also visualized the percent signal reduction at the consensus and expression-sensitive peaks present in both cell lines.

### Cell-cycle phased RNA-seq data processing

We downloaded the study’s sequencing files from NCBI GEO, then aligned the reads to the genome and transcriptome with STAR (v.2.7),^78^ allowing up to 20 multi-map reads and no more than 4% alignment mismatch. RSEM (v.1.3.3) was then used to quantify expression, specifying strandness as forward only.^79^ Then, for each of the three cell-cycle phases, G0/G1, S, and M, we took the quantified expression in FPKM for protein-coding genes. We performed similar upper-quartile normalization and converted the normalized expression levels to the log_2_ scale after adding a pseudocount of 1. We then visualized CTCF, SMC3, and RAD21 expression across the three phases.

Finally, we calculated Spearman correlation between CTCF occupancy and the average expression for genes upregulated in M-phase. We defined M-phase upregulation as at least 1.5-fold higher expression in M phase than in S phase and in G0/G1 phase, with at least 10 FPKM in M phase. This definition identified 1.6K genes to be upregulated in M phase. We computed the average expression of these genes across matched ENCODE samples and correlated it with CTCF occupancy.

### Promoter-to-TSS annotation

We downloaded refGene.txt from the UCSC hg38 database and only retained genes on the major chromosomes. These genes’ TSSs were then extracted, which were the start position for plus-strand genes and the end position for minus-strand genes. BedTools `closest` was used to associate the 126K consensus CTCF peaks with the closest TSSs.^80^ Peaks whose centers were within 2 kb of any TSS were considered TSS-proximal.

We then mapped promoters to TSSs. Similar to CTCF consensus peak list construction, we pooled the most 15,000 accessible peaks called from more than 1000 ENCODE DNase-seq and ATAC-seq datasets. These peaks were stitched into regions if they overlapped by at least 149 bp. For each region, Gaussian Mixture Models with one, two, and three components were fitted to all peak centers within the region. Regions were subdivided into one, two, or three regions, depending on which GMM had the lowest error. For each subdivided region, we formed a 300-bp peak centered on the region midpoint. Across the 973K accessible regions, 73K of them were proximal (<2 kb) to a TSS. Finally, we defined the region closest to the TSS as the gene promoter.

### GO analysis for genes proximal to TSS-proximal clusters

We used GSEApy to access the Molecular Signatures Database and retrieved the 2025 versions of c2 curated gene sets, c5 ontology gene sets, and hallmark gene sets, totaling 23839 gene sets. For each TSS-proximal cluster, we aggregated and deduplicated genes whose TSSs were proximal to peaks. Genes that were more than 2 kb from peaks in TSS-proximal clusters were excluded. GSEApy’s function `enrich` was used to perform hypergeometric tests. Based on the outputs, we subsequently calculated the log2 odds ratio and term ratio, defined as the ratio of overlapping genes to all genes in a gene set. We filtered out gene set entries with fewer than 30 overlapping genes, FDR greater than 10^−6^, and odds ratio smaller than 1. Then, for each peak cluster’s remaining gene sets, we sorted them by their log_2_ odds ratio and selected the top 10 for visualization.

### CpG content across CTCF peak clusters

For each peak cluster, we centered peaks on the strongest, STORM-called CTCF binding sites. Based on STORM-reported motif orientation, we reverse-complemented sequences whose motif was on the minus strand. For each cluster, we aligned all peak sequences extended to ±500 bp centered on the motif and calculated the percent of CpG dinucleotides at each position across all sequences. Global CpG background was computed as the ratio of the total number of CpG dinucleotides over all dinucleotides.

### CpG content stratified by CTCF peaks, Promoters, and TSSs

As described in the previous section, we associated consensus CTCF peaks with their closest TSSs. We also constructed a high-quality list of promoter-to-TSS mapping. We then built triplets of TSSs, promoters, and CTCF peaks by merging the two lists on TSSs. These triplets included CTCF peaks not in the TSS-proximal peak clusters. For each triplet, we computed distances between CTCF motifs, promoters, and TSSs. CTCF motifs and promoters were represented by their centers for the distance calculation.

For the first CpG analysis, we first separated the triplets based on the strand orientations of the associated genes. Then, for the plus-strand genes, we stratified triplets by whether CTCF motifs are upstream of promoters. For each stratum, we centered sequences on the center base pair of the CTCF motif. Sequences containing minus-orientation CTCF motifs were not reverse-complemented. CpG content was computed across aligned sequences. For the minus-strand genes, we similarly stratified triplets by whether motifs are upstream or downstream of promoters and calculated CpG content. For visualization, we performed a rolling average, with a window size of 9, except at the 19-bp CTCF motif.

For the second CpG analysis, we stratified triplets by relative ordering of CTCF motifs, promoters, and TSSs. Triplets containing minus-strand genes were merged with those containing plus-strand genes with reversed triplet order (e.g., motif-promoter-TSS was treated as TSS-promoter-motif). Then, for each triplet ordering, we aligned the three elements and rescaled the CpG content. We encoded all sequences as boolean vectors and extended each segment of the boolean vector to 10,000 values long. We then averaged the scaled vectors across all sequences within that stratum. No rolling average was applied here.

### Whole-Genome Bisulfite sequencing data processing

We downloaded WGBS-measured methylation state at CpG files associated with samples for which we had CTCF ChIP-seq from ENCODE (**Supp. Table 6**). As with ENCODE RNA-seq data, we additionally required sample donor information to match between CTCF ChIP-seq and WGBS. We then pooled CpG dinucleotide coverage (number of reads) and methylated reads across all datasets. When measurements were on the minus strand, we merged them with the plus-strand entries. We only kept dinucleotide entries on the major chromosome. We then built a methylation table, with rows representing dinucleotides for which at least one dataset had read coverage and columns representing the experiments. We pruned this table to only keep high-quality bases, defined as dinucleotides with at least 3 reads in at least five experiments. Then, we used BSmooth, with default parameters, to compute a smoothed methylation level across all remaining dinucleotides.^81^ We then intersected the dinucleotides with our consensus peaks using BedTools `intersect`, and for each peak we computed its methylation content as the average methylation level across all dinucleotides within it. We ended up having 117K peaks with a methylation readout. Methylation from datasets characterizing the same sample was averaged.

### Correlation between CTCF occupancy and DNA methylation level

We matched WGBS-measured methylation content across peaks with ChIP-seq-measured CTCF occupancy, by both sample and donor identities. Then Pearson correlation was computed for all peaks across 10 matching samples. Peaks with no methylation readout received a correlation of 0. We considered peaks to be CpG-enriched if they had at least 10 CpG dinucleotides. For each peak, we calculated methylation dispersion as the ratio between the variance of methylation across samples and the average methylation across samples. We then considered peaks methylation-variable if their dispersion was in the top 75th percentile, meaning only 25% of peaks were methylation-stable. Finally, only 14K peaks satisfied both criteria, CpG-enriched and methylation-variable. Among these peaks, we reported the distribution of Pearson correlation, stratified by whether the peak belongs to the TSS-proximal C3 cluster.

### Change in DNA methylation post TET triplet knockout

We downloaded ESC TET TKO (triple knockout TET1, TET2, TET3) per-CpG-dinucleotide methylation from NCBI GEO. The downloaded methylation files already filtered out all dinucleotides with fewer than 3 read coverages. The LiftOver tool was used to lift methylation information from hg19 to hg38. As in our WGBS data processing step, we intersected the dinucleotides with our consensus peaks using BedTools `intersect` and averaged methylation levels across all overlapping dinucleotides to represent per-peak methylation. We further removed peaks capturing fewer than three CpG dinucleotides. We then computed the change in methylation level between TET TKO and wild-type. Finally, we visualized the average methylation change across the remaining peaks for each cluster.

### Change in CTCF occupancy post DNMT1 interference

We downloaded K562 DNMT1i (DNA methyltransferase 1 interference) peak calls and signal tracks from NCBI GEO. BedTools `intersect` was used to identify consensus peaks that were occupied in the K562 cell line both in wild-type and in DNMT1 knockdown. BedTools `intersect` was also used for identification of peaks that were reactivated upon methylation removal. For each of the three peak groups, we extracted the average CTCF occupancy in both conditions across replicates. For each cluster, we calculated the fraction of peaks within it that became reactivated after DNMT1 knockdown and the fold change in CTCF occupancy among reactivated peaks.

### Cancer-cell line copy number variation analysis

Cancer cell-line copy number variation profiles (OmicsCNSegmentsWGS.csv) were downloaded from the DepMap portal under the 26Q1 release (https://depmap.org/portal).^82^ We then retrieved the copy number segment information inferred from the whole-genome sequencing data. Across our consensus peak list, we extracted the relative copy number in K562, HepG2, and MCF7 cell lines. Finally, we visualized the distribution of copy number ratios across constitutive peaks and cancer cell-line-enriched peaks in each of the three mentioned cancer cell lines.

### gkm-SVM and gkmPWM motif extraction procedure

To maximize the sensitivity of our motif extraction pipeline, we used only the peaks most confidently assigned to each cluster. So, for all peaks, we examined their nearest neighbors’ cluster membership. We kept those that shared a cluster membership with more than 75% of their neighbors. We focused on clusters that were not cancer cell-line-specific and had more than 300 peak sequences. From the remaining clusters, we trained pairwise gkm-SVM models.^20–22,52,83^ For each pair of clusters, we downsampled the larger cluster to match the smaller cluster in size, ensuring the two sets had the same distribution of sequence GC content. Because clustering is imperfect, subsampling also selected for peaks with dissimilar binding profiles.

gkm-SVM models were trained on subsampled sequences from each pair of clusters. Predictive power was evaluated with five-fold cross-validated auROC. To prevent data leakage, we split folds along major chromosomes.^84,85^ The first fold contains sequences on chr1, chr3, and chr6. The second fold contains sequences on chr2, chr8, chr9, and chr16. The third fold has chr4, chr11, chr12, chr15, and chrY sequences. The fourth fold has chr5, chr10, chr14, chr18, chr20, chr22 sequences. The last fold has peaks on chr7, chr13, chr17, chr19, chr21, and chrX.^64^ The division was chosen to ensure each fold has roughly the same total chromosome length. This cross-validation procedure yielded one file of predicted scores for performance evaluation and one file of the gkm-SVM model for motif extraction.

We ran gkmPWMlasso (v.1.0) with default parameters on all pairwise gkm-SVM models to extract motifs.^49^ We supplied it with the JASPAR 2024 core vertebrate redundant motif list and asked for 30 motifs.^28^ Thus, we obtained a gkmPWMlasso motif call for each trained model. We then tabulated all mentioned motifs and filtered for non-redundant ones. To identify the non-redundant motifs, we clustered the 1912 motifs in the JASPAR 2024 database by Pearson similarity. We used the graph-based algorithm to identify the non-redundant motif sets. To do so, we represented each motif as a node in the graph, and we added edges between all motif pairs with correlation coefficients above 0.93. Finally, we extracted all connected components from the graph, and each connected component represented a non-redundant motif type.

Then, for each motif type, we retained those that occurred more than 10 times across 300+ gkmPWMlasso reports and had an average absolute gkmPWMlasso-reported z-score greater than 1.8. This resulted in around 60 non-redundant, informative motifs. We then combined pairwise gkm-SVM models that had the same positive class to create quasi one-vs-all model files. gkmPWMlasso was rerun on these one-vs-all models against the non-redundant, informative motifs to re-estimate the frequency and statistical significance of these motifs across all clusters.

### Consistency between motif scanning and gkmPWM motif extraction

We converted the 60 non-redundant, informative motifs from MEME format to STORM-compatible format. STORM assigned motif scores for all supplied motifs to all CTCF peak sequences. For each motif, we binned peaks into 20 bins based on the associated motif score, from the weakest to the strongest. Then, for each cluster, we computed the ratio of the number of peaks in the second-highest bin to the number in the second-lowest bin. We called this ratio motif enrichment. Then, we computed the Spearman correlation between motif enrichment and gkmPWMlasso-reported z-score across all clusters.

### Cell-type-specific CTCF peaks and enhancer k-mer correlation

For all clusters with one-vs-all models, we computed the predicted scores for all possible length-11 k-mer (https://github.com/Dongwon-Lee/lsgkm#generating-weight-files-for-deltasvm). As described in the previous literature, we scored all length-11 k-mer with tissue- and cell-type-specific enhancer models.^49^ We then correlated the scores for 11-mers generated from the cell-type-specific CTCF peaks and enhancer peaks. High correlation was observed among CTCF models and enhancer models that had the matching cell and tissue types.

### Characterization of CTCF-peak-overlapping dsQTLs

We extended the 574 lymphoblastoid-cell-line dsQTLs to 300-bp peaks in hg38.^64,65^ BedTools `intersect` was used to overlap dsQTL peaks and consensus CTCF peaks, requiring at least 150-bp overlap. This meant that 220 dsQTL base pairs fell within the 300-bp CTCF peaks. Then, we chose a set of 14 TF motifs predicted to be highly relevant in GM12878, a lymphoblastoid cell line (https://beerlab.org/gkmpwm/pdfs/DHS_553_gkmPWMlasso.pdf). The TFs were EBF4, IKZF1, RUNX2, NFKB, SPI1, IRF3, MEF2, AP1, FERD3L, NRF1, NFY, SP1, YY1, and THAP11. We scored the dsQTL-flanked sequences for each TF motif, including CTCF, with STORM. The sequences were long enough to accommodate all possible motif-scoring windows that contained the dsQTL base pair. Both the reference and the alternative allele versions of the 220 dsQTL sequences were scored by all 15 TFs. For each sequence, we called it to disrupt/gain the motif with the largest change in motif score between the two alleles. Motif scores were capped at 0, which indicated a random chance of observing the TF in that sequence. Finally, we tabulated the fraction of CTCF-peak-overlapping dsQTLs that alter CTCF binding sites compared to the other GM12878-specific TF binding sites.

### Reduced peak list analysis

We used the same set of high-quality CTCF ChIP-seq experiments to generate the reduced consensus peak list. Similar to before, all peaks reaching the statistical threshold were stitched into regions, and only regions supported by at least three experiments were retained. We still required the region-center-derived peaks not to overlap the ENCODE blacklist or have a CTCF motif score above 0. However, rather than requiring the peak signal to be at least 25% of the 3000th-strongest peak’s signal, we tightened the threshold to be at least 40% of the 3000th-strongest peak’s signal. This stricter filtering reduced the number of consensus peaks from 126K to 82K. BedTools `intersect` was used to lift the previous cluster membership assignment to the reduced list. This allowed us to precisely calculate the number of peaks retained for each category. All downstream analyses, including correlating CTCF occupancy to CTCF expression, distribution of GC content and distance to the nearest TSS across clusters, and gkm-SVM predictive performance, were the same as those done on the full consensus peak list.

### Histone ChIP-seq data processing

H3K4me1, H3K4me3, and H3K27me3 ChIP-seq were obtained from the ENCODE consortium (encodeproject.org). We downloaded datasets associated with the ten biological samples for which we have CTCF ChIP-seq and WGBS data (**Supp. Table 7**). Of the ten samples, five were cell lines, and five were tissue samples, such as the heart left ventricle and the upper lobe of the left lung. As described previously for CTCF ChIP-seq, we used MACS2 for peak calling and signal track generation. Then, for each dataset, we extracted the average signal within a 2000 bp window centered on each peak midpoint. For each histone modification, we computed a normalized scaling factor for each dataset to achieve mean normalization.

Then, we used BedTools `random` and `subtract` to obtain a list of randomly sampled 300-bp windows that did not overlap with our consensus CTCF peak list. We computed these background regions’ GC content and scored them for CTCF motif strength with STORM. For each peak cluster, we subsampled an equal-sized, GC-matched, CTCF motif-score-matched set from the background regions. We sampled background regions for fibroblast-specific, LCL-specific, K562-specific, HepG2-specific, heart-specific, and TSS-proximal C1/C2 clusters. We finally extracted average signals from 2000 bp windows centered on peak midpoints and background-region midpoints. These signals were normalized with the previously calculated normalization factor, and then we visualized the signals at the peaks versus those at background regions.

### Regression model for CTCF occupancy

To predict quantitative CTCF occupancy at our consensus peaks, we prepared seven features for ten biological samples. CTCF expression was the only feature that was unique to each sample. Three features were unique to each peak, independent of sample: CTCF motif scores, GC content, and the proximal genes’ average expression across samples. The last three features were specific to each combination of peak and sample, and they were H3K4me1 signal, H3K27me3 signal, and WGBS-measured methylation level. In the previous sections, we have described how we obtained these features. To prevent data leakage, the proximal genes’ expression was averaged across samples other than the ten that the regression task was structured around. For most of the CTCF peaks without a proximal gene, this field was 0. Meanwhile, the methylation level was defined as the change in methylation from the mean methylation averaged across samples outside the ten samples.

Therefore, the feature matrix had ten times the peak count of rows and seven columns representing features. The observed CTCF occupancy was transformed to the log_2_ scale for variance stabilization. We used a shallow XGBoost regressor (v3.2), with mostly default parameters (n_estimators=500, learning_rate=0.05, min_child_weight=10, subsample=0.8, colsample_bytree=0.8), to predict CTCF occupancy from the seven features. Predictive performance was not sensitive to parameter choices, except for tree depth. We robustly evaluated regressor performance using a leave-one-sample-out scheme, so the regressor had never seen the test set’s feature space during training.

### Shapley contribution

Per our leave-one-sample-out cross-validation scheme, we obtained all predicted occupancy after training ten separate XGBoost regressors. We evaluated the model performance in three different ways. We first checked how well the model predicted occupancy. Thus, we correlated predicted and observed signals across peaks, stratified by either samples or peak categories. For the sample stratification, ten Pearson correlation coefficients were computed. In addition, for each peak, we correlated predicted and observed signals across the ten biological samples. The hundred thousand correlation values were then organized by peak category.

We next investigated how each feature contributed (a.k.a. Shapley contribution) to across-peak and across-sample correlations for each peak category. To do so, we trained the XGBoost regressor on all possible feature combinations. Because we had seven features, there were 127 combinations of features (127=2^7^-1). Then, for each peak category and each feature of interest, we computed the change in correlation with and without the feature (e.g., correlations from a regressor trained on the CTCF motif score and H3K4me1 versus correlations from a regressor trained on the CTCF motif score alone). When excluding the feature of interest, we were left with six features, thus, there were 64 feature combinations (64=2^6^), meaning 64 calculation of delta correlations. For the final Shapley value, for each peak category and each feature of interest, we average the delta correlation across these 64 correlation coefficients.

### Hi-C data processing

We obtained Hi-C mapping-quality-thresholded contact matrices from the ENCODE consortium (encodeproject.org). We downloaded datasets for which we had CTCF ChIP-seq data. We additionally required sample donor information to match between CTCF ChIP-seq and Hi-C (**Supp. Table 8**). We ended up with 40 distinct biological samples. For each sample, we selected the dataset with the highest read depth. Then, from the downloaded contact map, we extracted the 5 kb-resolution matrices and converted them to the cool format with hictk.^86^ We normalized the 5 kb matrices using cooler `balance`, with default parameters.^87^ The procedure also marked some bins as unreliable, for reasons like low read depth.

### Insulation score

Cooltools `insulation` was used to calculate insulation on 100 kb windows.^88^ For each distinct biological sample, we retrieved the insulation scores calculated from the corresponding Hi-C matrix for peaks that were present. For this analysis, we considered a peak present in a biological sample if it had sufficiently high signal across all datasets that characterized this sample. We also dropped insulation scores for peaks that were in an unreliable bin (described in the Hi-C data processing section). Insulation scores were standardized (mean=0, standard deviation=1). Then, for each biological sample and each peak category, we reported the mean insulation scores.

### Contact frequency and Occupancy

Starting with the weighted, 5 kb-resolution Hi-C matrices stored in .cool format, we first computed expected counts by distance. For each matrix, we averaged the contact frequency for all pairs of contacts that were separated by the same distance. We kept only pairs with both bins deemed reliable by cooler, and that were both on the major chromosomes. This yielded an expected frequency vector for all distances between 5 kb and 1 Mb, for each dataset.

Starting with the consensus peak list, we identified 6.7 million putative pairs of CTCF peaks that were further than 5 kb and within 1 Mb of each other. We extracted the contact frequency for all candidate pairs across all datasets. We divided the contact frequencies extracted from matrices by the corresponding expected frequencies, depending on the distances between the pairs. Using the contact-frequency matrix, we identified some experiments as likely poor quality because they showed very low correlation with all other experiments. A few more datasets were removed because the matrix size was much smaller than others.

Among the remaining 33 datasets, we filtered putative CTCF peak pairs based on whether their motifs were in convergent orientation. Convergent orientation meant that the left anchor contained a STORM-called binding site on the plus strand and the right anchor contained a minus-strand binding site. We further required the peak pair to have a contact frequency of at least seven in at least one experiment, and for both peaks to be present in at least one associated biological sample. For this analysis, we considered a peak present in a sample if it had sufficiently high signal in at least half of the datasets characterizing the sample. We identified 25K interacting peak pairs.

Meanwhile, we sampled an equal number of non-interacting pairs from peaks present in at least one sample. We defined the pairs with contact frequency below two in all surveyed Hi-C matrices. Finally, for these two sets of peaks, we computed the Pearson correlation between contact frequency and CTCF occupancy. Occupancy was defined as the sum of CTCF occupancy at both anchors, averaged across all datasets characterizing each of the biological samples. We compared the distributions of correlation coefficients for interacting pairs and non-interacting pairs and visualized the distribution for loops anchored by specific peak categories. For instance, we examined pairs where both anchors were constitutive peaks or those where one anchor was constitutive and the other was expression-sensitive.

### ChIA-PET data processing

We obtained ChIA-PET read pairs in BEDPE format from the ENCODE consortium (encodeproject.org), two representing fibroblasts and two representing lymphoblastoid cell lines (**Supp. Table 9**). We kept only the read pairs where both reads were on the same chromosome, within 1 Mb of each other, and where both anchors’ start and end positions were within 5 kb. We then mapped the left and right anchors of each read pair to their nearest consensus CTCF peaks. Only pairs with both anchors within 2 kb of CTCF peaks were kept. This allowed us to extract interaction strengths between pairs of consensus CTCF peaks in fibroblasts and LCLs. Loop interaction strength among all detected peak pairs was normalized to a sum of 1e6. We then calculated the log_2_ fold change in loop strength between fibroblasts and LCL samples and visualized the average log fold change among a subset of peaks stratified by anchor cluster membership. For instance, loops where both anchors were fibroblast-specific, or those where the left anchor was in the LCL-specific cluster and the right anchor was in the expression-sensitive cluster.

For each gene, we summed the strength of all loops that enclosed the gene’s TSS. Loop enclosure was defined as TSSs being located between the left and the right anchors of a loop. We then calculated the log_2_ fold change in the strength of loops enclosing TSSs. We then ranked the protein-coding genes by their log_2_ fold change in gene expression in fibroblasts versus in LCL. The top 1000 upregulated and top 1000 downregulated genes in fibroblasts were then identified. We finally predicted whether these 2000 genes were up- or down-regulated based on whether the loops enclosing them. We also recomputed the log_2_ fold change in loop strength using only loops with both anchors in the constitutive cluster and evaluated loop strength as a predictor of the direction of differential expression. For the fibroblast-specific-only, LCL-specific-only, or expression-sensitive-only sets, we recomputed the change in loop strength with loops with at least one anchor belonging to these clusters.

### Visualization

All data visualization in this manuscript was performed with Matplotlib and Seaborn.^89,90^

